# Isogenic sets of hiPSC-CMs harboring *KCNH2* mutations capture location-related phenotypic differences

**DOI:** 10.1101/846519

**Authors:** Karina O. Brandão, Lettine van den Brink, Duncan C. Miller, Catarina Grandela, Berend J. van Meer, Mervyn P.H. Mol, Leon G.J. Tertoolen, Christine L. Mummery, Luca Sala, Arie O. Verkerk, Richard P. Davis

## Abstract

**Aims:** Long QT syndrome type 2 (LQT2) is caused by mutations in the gene *KCNH2*, encoding the hERG ion channel. Clinically, mild and severe phenotypes are associated with this cardiac channelopathy, complicating efforts to predict patient risk. The location of the mutation within *KCNH2* contributes to this variable disease manifestation. Here we determined whether such phenotypic differences could be detected in cardiomyocytes derived from isogenic human induced pluripotent stem cells (hiPSCs) genetically edited to harbour a range of *KCNH2* mutations.

**Methods and Results:** The hiPSC lines heterozygous for missense mutations either within the pore or tail region of the ion channel were generated using CRISPR-Cas9 editing and subsequently differentiated to cardiomyocytes (hiPSC-CMs) for functional assessment. Electrophysiological analysis confirmed the mutations prolonged the action potentials and field potentials of the hiPSC-CMs, with differences detected between the pore and tail region mutations when measured as paced 2D monolayers. This was also reflected in the cytosolic Ca^2+^ transients and contraction kinetics of the different lines. Pharmacological blocking of the hERG channel in the hiPSC-CMs also revealed that mutations in the pore-loop region conferred a greater susceptibility to arrhythmic events.

**Conclusion:** These findings establish that subtle phenotypic differences related to the location of the *KCNH2* mutation in LQT2 patients are reflected in hiPSC-CMs under genetically controlled conditions. Moreover, the results validate hiPSC-CMs as a strong candidate for evaluating the underlying severity of individual *KCNH2* mutations in humans which could ultimately facilitate patient risk stratification.

**Translational perspective:** Clinical management of patients diagnosed with cardiac channelopathy diseases such as LQT2 is complicated by the variable disease phenotypes observed among mutation carriers, creating challenges for diagnosis, risk stratification and treatment. The genotype of the patient contributes to this clinical heterogeneity, with the influence of the mutation’s location within *KCNH2* on a patient’s risk of a cardiac event being an example. Here we demonstrate that under stringently controlled genetic and experimental conditions, hiPSC-CMs are able to reflect these subtle genotype-phenotype differences, thereby providing new opportunities to stratify and potentially lessen sudden cardiac death risk amongst *KCNH2* mutation carriers.

## 1. Introduction

Congenital long QT syndrome (LQTS) is a genetic disease with an estimated prevalence of ∼1:2000 individuals. It is characterised by a prolonged QT interval on an electrocardiogram that can lead to sudden cardiac death, particularly in young people ^1,2^. Although the identification of genes associated with LQTS has dramatically improved our understanding of the disease ^3^, clinical management remains complicated by the variability in disease expressivity and penetrance among mutation carriers which range from lifelong asymptomatic to experiencing life-threatening arrhythmias ^4^. While environmental factors are contributors to this clinical heterogeneity ^5^, genetics also plays a major role through both the primary genetic mutation and the presence of additional genetic variants that modify the disease outcome ^6,7^.

Type 2 LQTS (LQT2) is the second most prevalent form of congenital LQTS and is due to mutations in *KCNH2*, which encodes the α subunit of the Kv11.1 (hERG) channel responsible for conducting the rapid delayed rectifier potassium current (I_Kr_) in cardiomyocytes ^8^. Several studies have demonstrated that the location of the mutation within this ion channel is an important determinant of arrhythmic risk in LQT2 patients, with patients harbouring mutations in the pore-loop region at higher risk of cardiac events than those with mutations located in other regions ^9–12^. Furthermore, mutations that result in a dominant negative effect, in which the function of wildtype hERG is reduced or eliminated, also produce higher adverse event rates ^12,13^. However mutations identified within the pore-loop region can, in some instances, result in a less severe outcome ^14^, highlighting the need for *in vitro* models to accurately classify these rare variants and for gaining mechanistic insights into their contribution to disease phenotypes.

Cardiomyocytes derived from human induced pluripotent stem cells (hiPSCs) are now well-established as models for LQT2 ^15–18^. Indeed, a number of hiPSC lines have been derived from both symptomatic and asymptomatic patients with mutations in various regions of hERG ^19^. However, as these lines are from different individuals, they harbour additional genetic variants that may functionally influence the disease phenotype observed and limit the utility of hiPSC-derived cardiomyocytes (hiPSC-CMs) for broad intragenotype risk stratification.

To create a tailored model to study the genetic aetiology of LQT2, we generated a set of isogenic hiPSC lines that possess heterozygous mutations within the pore-loop domain (KCNH2-A561T) or in the cytoplasmic tail (KCNH2-N996I) of hERG by genetically modifying a control hiPSC line. Molecular and functional comparisons of these edited lines confirmed not only that the *KCNH2* variant hiPSC-CMs phenocopied the key features of LQT2 but also that differences due to the mutation were reflected in the cell lines. This included dissimilarities in the mechanism underlying the hERG channel trafficking defect caused by these mutations, as well as a more prolonged repolarisation observed in hiPSC-CMs with the pore mutation when measured as a paced syncytium. Furthermore, when these hiPSC-CMs were exposed to E-4031, a hERG channel blocker, they were more susceptible to proarrhythmic effects compared to either the hiPSC-CMs with the tail mutation or the unedited control. Our findings demonstrate the potential of hiPSC-CMs to reveal the inherent severity of individual *KCNH2* mutations when using genetically-matched lines but also further advances hiPSC-CMs as a robust model for not only predicting risk but also in the development of better risk stratification for patients.

## 2. Methods

An extended methods section is provided in the Supplementary material.

### 2.1 Genome editing

The *KCNH2* variants (c.G1681A and c.A2987T) were introduced into a hiPSC control (KCNH2^WT/WT^; LUMC0020iCTRL-06) ^20^ line by CRISPR/Cas9-mediated gene editing. Heterozygosity of clones was confirmed by Sanger sequencing. Sequences of the guide RNAs (gRNAs), single-strand oligonucleotides (ssODNs), PCR primers and the resulting hiPSC lines used in this study are listed in Supplementary material tables S1-S3.

### 2.2 Differentiation to hiPSC-CM

The hiPSC lines were differentiated into cardiomyocytes as described in the Supplementary material. All analyses were performed on cryopreserved hiPSC-CMs 5-9 days after thawing.

### 2.3 Electrophysiology

Voltage-clamp recordings of I_Kr_ were made using pipette and bath solutions as previously described ^17^. Action potentials (APs) of the hiPSC-CMs were recorded by perforated patch-clamp and the dynamic clamp technique with injection of an inward rectifier potassium current (I_K1_) used to achieve a close-to-physiological resting membrane potential (RMP) ^21^. For electrophysiological analysis on multi-electrode arrays (MEAs), the field potential (FP) was recorded as previously described ^22^. Sequential addition of increasing concentrations of E-4031, a specific I_Kr_ blocker, was performed with recordings initiated following 1 min of incubation.

### 2.4 Optical recordings

The hiPSC-CMs were labelled with organic fluorescent dyes and the resulting signals recorded and analysed as described in the Supplementary material.

### 2.5 Statistical Analysis

Results are presented as mean ± SEM, with comparison between groups performed using one-way or two-way ANOVA followed by Tukey’s multiple comparisons test for post hoc analysis. Pairwise comparisons were additionally performed using the Student’s t-test following one-way ANOVA if one of the null hypotheses could be rejected ^23^. Curve fitting from regression models and statistical analyses were performed with Graphpad Prism 8 software, with a *P* value <0.05 considered statistically significant.

## 3. Results

### 3.1 Generation and characterisation of an isogenic set of KCNH2 variant hiPSC lines

A limitation of hiPSC lines derived from unrelated patients with different LQT2-causing mutations is the inability to compare the resulting hiPSC-CMs under genetically-matched conditions. Therefore, to detect phenotypic differences arising from the region in which the mutation was located in *KCNH2*, we elected to genetically introduce these mutations into a well-characterised hiPSC line derived from a healthy individual (KCNH2^WT/WT^) ^20^. Furthermore, we confirmed that this cell line did not carry any known disease-causing mutations by performing whole exome sequencing and examining a panel of 107 genes known to be linked to inherited arrhythmia syndromes or cardiomyopathies ^24^ (Supplementary Table S4). All coding sequence variants identified were predicted to be benign due to their frequency in the general population being ≥1%. The only exception was a rare variant identified in *DOLK*, the gene encoding dolichol kinase. Homozygous mutations in this gene can lead to multi-systemic glycosylation disorders including dilated cardiomyopathy, with individuals typically not surviving to adulthood ^25^. However, the variant identified in the KCNH2^WT/WT^ line is unlikely to be pathogenic as the hiPSCs were derived from a healthy 51-year old female and were heterozygous for the *DOLK* variant.

We then used a CRISPR/Cas9-mediated gene editing strategy to generate an isogenic set of hiPSC lines harbouring a missense variant either within the pore-loop domain (NM_000238.3:c.1681G>A, NP_000229.1:p.Ala561Thr) or cytoplasmic tail region (NM_000238.3:c.2987A>T, NP_000229.1:p.Asp996Iso) of *KCNH2* (Figure 1). RFLP analysis identified clones that appeared to be genetically modified (Supplementary Figure S1) and these were subsequently confirmed by Sanger sequencing to be either heterozygous for the KCNH2-A561T (KCNH2^PR/WT^; Figure 1E) or the KCNH2-N996I (KCNH2^TL/WT^; Figure 1F) variant. For each mutation, a second independent isogenic heterozygous clone (KCNH2^PR/WT^ cl2 and KCNH2^TL/WT^ cl2) was also selected for further characterisation (Supplementary Figure S2 and S3). All clones were assessed by Sanger sequencing for potential off-target modifications due to the CRISPR/Cas9 transfection, with no insertions or deletions detected at any of the genomic loci examined (Supplementary Table S5). Furthermore, G-band karyotyping of these clones indicated the lines were karyotypically normal, and the undifferentiated hPSCs expressed the stem cell markers SOX2, OCT-4, SSEA4, NANOG (Supplementary Figure S2-S4).

**Figure 1.**
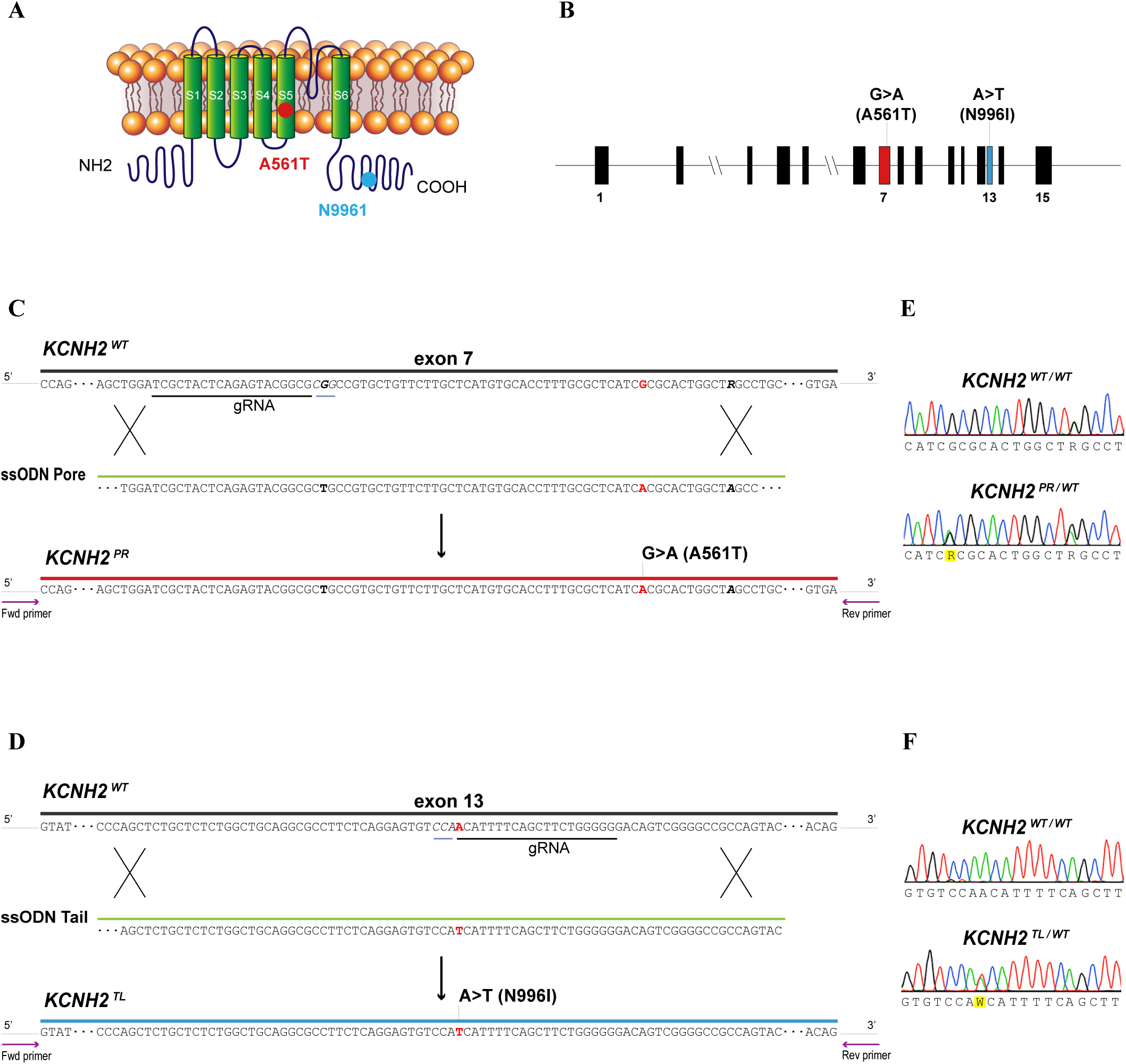
Generation of isogenic hiPSC lines with *KCNH2* mutations. **(A)** Structure of the potassium ion channel hERG encoded by *KCNH2* indicating the introduced mutations (KCNH2-A561T, red dot; KCNH2-N996I, blue dot). **(B)** Structure of the *KCNH2* genomic locus highlighting the exons modified to generate the KCNH2 variant hiPSC lines. **(C, D)** Schematic outlining strategy to introduce the KCNH2-A561T mutation (*KCNH2^PR^*) **(C)** or the KCNH2-N996I mutation (*KCNH2^TL^*) **(D)** by homologous recombination into a *KCNH2* wildtype (*KCNH2^WT^*) sequence. The gRNA and their corresponding protospacer adjacent motif sequences are underlined in black and grey respectively. Part of the ssODN sequences to introduce the A561T (ssODN Pore) and N996I (ssODN Tail) mutations are shown. Nucleotides modified to introduce the mutations are indicated in red, silent mutations and SNPs used to assist with the targeting and screening are bolded in black. Arrows represent the PCR primers used to identify targeted clones. **(E, F)** Sequence analysis of the PCR-amplified genomic DNA showing heterozygous introduction of NM_000238.3:c.G1681A **(E)** and NM_000238.3:c.A2987T **(F)**.

### 3.2 Mutations in distinct regions of hERG cause differing trafficking defect phenotypes

A monolayer-based differentiation protocol was used to differentiate the KCNH2 variant and KCNH2^WT/WT^ hiPSC lines to cardiomyocytes. Flow cytometric analysis for the pan-cardiomyocyte marker cardiac troponin T (cTnT) indicated that all lines, except KCNH2^PR/WT^ cl2, differentiated with similar efficiencies, with on average ∼70-80% of the cells being cTnT^+^ (Figure 2A and B; Supplementary Figure S5). For functional comparisons between the cell lines it is important that the cardiomyocyte population contains a similar proportion of ventricular cells. The hiPSC-CMs (cTnT^+^ cells) were therefore further characterised for the expression of the ventricular cardiomyocyte marker, MLC2v. On average, the proportion of hiPSC-CMs that were ventricular was between 45-64%, including for the KCNH2^PR/WT^ cl2 line, (Figure 2A and B; Supplementary Figure S5). The hiPSC-CMs also displayed characteristic sarcomeric structures that were positive for α-actinin and myosin heavy chain (Figure 2C).

**Figure 2.**
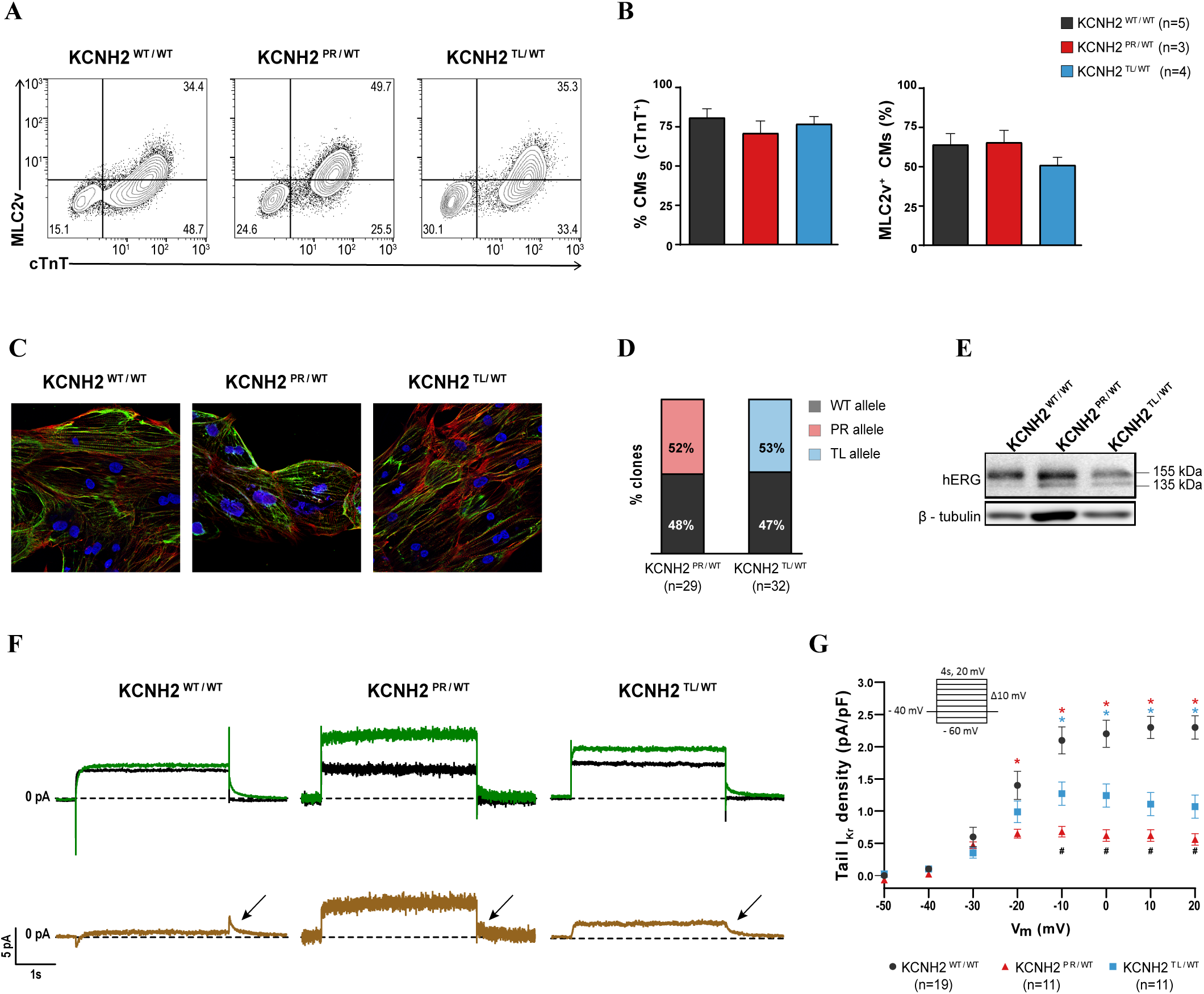
Evaluation of hERG channel function in differentiation day 21+7 KCNH2^WT/WT^, KCNH2^PR/WT^ and KCNH2^TL/WT^ hiPSC-CMs. **(A)** Representative flow cytometry plots of hiPSC-CMs for expression of cTnT and MLC2v in the indicated lines. Values inside the plots are the percentage of cells within the gated region. (**B)** Overall cardiac differentiation efficiency of the three hiPSC lines, showing average percentage of hiPSC-CMs (cTnT^+^) (left graph), and proportion of ventricular-like (MLC2v^+^) cardiomyocytes within the hiPSC-CM population (right graph). **(C)** Immunofluorescence images of the cardiac sarcomeric proteins α-actinin (red) and myosin heavy chain isoforms α and β (green) in the indicated lines. Nuclei (blue) were stained with DAPI (original magnification, 40x). **(D)** Percentage of *KCNH2* mRNA in the KCNH2^PR/WT^ and KCNH2^TL/WT^ hiPSC-CMs transcribed from the wildtype (WT) or mutated (PR, TL) alleles. **(E)** Western blot analysis of hERG in the indicated lines. Bands corresponding to core- and complex-glycosylated hERG (135 and 155 kDa, respectively) are marked. β-tubulin was used as a loading control. **(F)** Top panels show representative current traces evoked by a 4s voltage step from −40 to 0 mV before (green lines) and after (black lines) application of 5 µM E-4031. Bottom panels present the E-4031-sensitive current (brown lines). Arrows indicate the tail current of I_Kr_. **(G)** Average current-voltage relationships for I_Kr_ tail density in the indicated hiPSC-CMs. Inset: voltage protocol; * indicates statistical significance to KCNH2^WT/WT^ (*P*<0.0001); # indicates statistical significance between KCNH2^PR/WT^ and KCNH2^TL/WT^ (−10mV, 0mV *P*<0.001; 10mV, 20mV, *P*<0.01; two-way ANOVA with Tukey’s multiple comparisons for *post hoc* test).

Both KCNH2^PR/WT^ and KCNH2^TL/WT^ hiPSC-CMs expressed equal fractions of mutant and wildtype *KCNH2* transcripts (Figure 2D), confirming that the introduced mutations did not disrupt biallelic expression of the gene and suggesting that any differences observed between the cell lines would be due to dysfunction of hERG. Western blot analysis for hERG identified two protein bands – the fully glycosylated mature form (155 kDa) and the core-glycosylated precursor form (135 kDa). While in KCNH2^WT/WT^ hiPSC-CMs the mature form was predominantly present, in both KCNH2^PR/WT^ and KCNH2^TL/WT^ hiPSC-CMs higher expression of the precursor form was detected (Figure 2E), indicating that both mutations lead to a defect in trafficking of hERG to the cell membrane.

Finally, we determined how the trafficking defects caused by both *KCNH2* mutations affected I_Kr_. Representative examples of current traces in individual hiPSC-CMs from both KCNH2 variant and KCNH2^WT/WT^ hiPSC-CMs are shown in Figure 2F, with I_Kr_ measured as an E-4031-sensitive current. The I_Kr_ tail density was significantly reduced in both KCNH2^PR/WT^ and KCNH2^TL/WT^ hiPSC-CMs compared to the KCNH2^WT/WT^ hiPSC-CMs (Figure 2G). For example, after a voltage step to +10 mV, the I_Kr_ tail density was decreased by 51% in the KCNH2^TL/WT^ hiPSC-CMs and by 73% in the KCNH2^PR/WT^ hiPSC-CMs. Taken together, these findings suggest that although both *KCNH2* variants result in a trafficking defect of hERG, the KCNH2^TL/WT^ variant leads to haploinsufficiency while the KCNH2^PR/WT^ variant causes a dominant-negative phenotype.

### 3.3 The dominant-negative KCNH2 mutation causes a more severe electrical phenotype

To determine whether the differences in I_Kr_ density between the two KCNH2 variants was also reflected in the AP duration (APD), APs from individual hiPSC-CMs were recorded (Figure 3A). The APD at 50% and 90% repolarisation (APD_50_ and APD_90_, respectively) were significantly prolonged for both KCNH2 variant lines (KCNH2^PR/WT^: 185±17ms (APD_50_), 243±19ms (APD_90_); KCNH2^TL/WT^: 198±16ms (APD_50_), 254±18ms (APD_90_)) compared to the KCNH2^WT/WT^ hiPSC-CMs (131±11ms (APD_50_), 151±11ms (APD_90_)), however there was no significant difference between the two KCNH2 variant lines (Figure 3B). At both slower and faster pacing frequencies (0.2–4Hz), the differences in APD_90_ between the KCNH2 variant and wildtype lines remained, with no difference between the KCNH2^PR/WT^ and KCNH2^TL/WT^ hiPSC-CMs (Supplementary Figure S6). Additionally, arrhythmogenic activity as evidenced by the presence of early after depolarisations (EADs), during 0.2 Hz stimulation was detected in both KCNH2 variant lines but not the KCNH2^WT/WT^ line (Figure 3C and D). Also here, there was no difference in the frequency of EADs between the two lines (KCNH2^PR/WT^: 18.2%; KCNH2^TL/WT^: 17.6%). No significant differences in AP amplitude (APA) and RMP were observed between any of the lines, and while upstroke velocity (V_max_) appeared faster in the KCNH2^PR/WT^ hiPSC-CMs, this was not significant (*P*=0.07) (Figure 3B).

**Figure 3.**
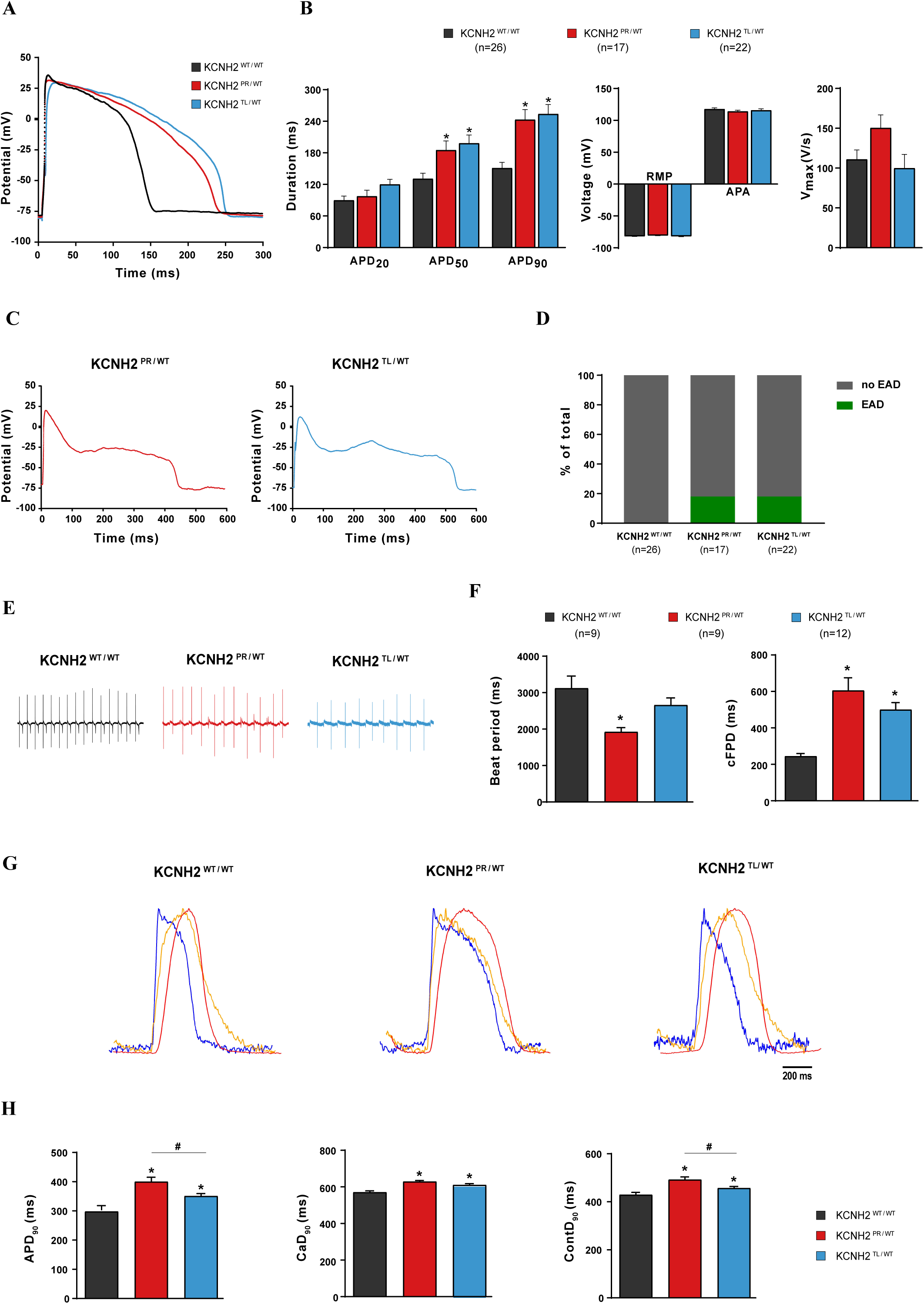
Electrophysiological characterisation of the KCNH2^WT/WT^, KCNH2^PR/WT^ and KCNH2^TL/WT^ hiPSC-CMs. **(A, B)** Representative AP traces **(A)**, and average APD_20,_ APD_50_, APD_90_, RMP, APA, and V_max_ values **(B)** for the indicated lines paced at 1 Hz. * indicates statistical significance to KCNH2^WT/WT^ (KCNH2^PR/WT^: APD_50,_ *P*<0.05; APD_90,_ *P*<0.001; KCNH2^TL/WT^: APD_50,_ *P*<0.01; APD_90,_ *P*=0.0001; one-way ANOVA with Tukey’s multiple comparisons for *post hoc* test). **(C, D)** AP traces for KCNH2^PR/WT^ and KCNH2^TL/WT^ hiPSC-CMs showing representative EADs **(C)**, and percentage of the indicated lines displaying EADs **(D)** when paced at 0.2 Hz. **(E, F)** Representative MEA traces **(E)**, and average values for beat period and cFPD **(F)** for the indicated lines. * indicates statistical significance to KCNH2^WT/WT^ (KCNH2^PR/WT^: beat period, *P*<0.01; cFPD, *P*<0.0001; KCNH2^TL/WT^: cFPD, *P*=0.001; one-way ANOVA with Tukey’s multiple comparisons for *post hoc* test). **(G)** Representative averaged timeplots of baseline-normalised fluorescence signals for the indicated lines stimulated at 1.2 Hz. AP traces are shown in blue, cytosolic Ca^2+^ flux in orange, and contraction-relaxation in red. **(H)** Average APD_90_, CaD_90_, and ContD_90_ values for the indicated lines as determined by changes in fluorescent signal. * indicates statistical significance to KCNH2^WT/WT^ (KCNH2^PR/WT^: APD_90_, CaD_90_, ContD_90_, *P*≤0.0001; KCNH2^TL/WT^: APD_90,_ ContD_90_, *P*<0.05; CaD_90_, *P*<0.001); # indicates statistical significance between KCNH2^PR/WT^ and KCNH2^TL/WT^ (APD_90_, ContD_90_, *P*<0.05); n=35-39 recordings for each cell line (Student’s t-test following one-way ANOVA with Tukey’s multiple comparisons for *post hoc* test).

Next, we investigated whether differences could be detected between the KCNH2^PR/WT^ and KCNH2^TL/WT^ hiPSC-CMs if measured in a syncytium. Figure 3E shows representative FP recordings of the hiPSC-CMs obtained from a MEA platform. Differences in beating frequency were observed between the 3 lines, with the KCNH2^PR/WT^ hiPSC-CMs showing a significantly faster beat period than the control KCNH2^WT/WT^ hiPSC-CMs (1920±123ms vs 3117±338ms). Therefore, the FPD was corrected (cFPD) for beat rate variability according to Bazett’s formula (Figure 3F). Mirroring the APD differences observed in single hiPSC-CMs, the cFPD was significantly prolonged in KCNH2^PR/WT^ and KCNH2^TL/WT^ hiPSC-CMs (605±70ms and 499±40ms, respectively) compared to KCNH2^WT/WT^ hiPSC-CMs (245±15ms), but was not significantly different between the two KCNH2 variant lines. The second KCNH2^PR/WT^ and KCNH2^TL/WT^ clones corroborated these findings with similar beat period and cFPD values (Supplementary Figure S7). Collectively, these data indicate that the genetically engineered KCNH2^PR/WT^ and KCNH2^TL/WT^ hiPSC-CMs displayed the expected electrophysiological characteristics of the LQT2 phenotype, however no differences in baseline measurements or frequency of arrhythmic events were found between the hiPSC-CMs harbouring these mutations.

Finally, we also evaluated the KCNH2 variant lines using a high-speed optical system that can simultaneously measure the AP, intracellular Ca^2+^ transients and contraction-relaxation kinetics of hiPSC-CM monolayers under paced conditions ^26^. This enables the rapid assessment of how LQT2-causing mutations affect the complete excitation-contraction coupling cascade. Representative transients of the three measured parameters for each of the lines are shown in Figure 3G. Analysis of the voltage traces also showed a significant increase in APD_90_ for both the KCNH2^PR/WT^ and KCNH2^TL/WT^ hiPSC-CMs (403±35ms and 350±21ms, respectively) compared to the KCNH2^WT/WT^ hiPSC-CMs (297±33.5ms; Figure 3H). Both Ca^2+^ transient and contraction at 90% duration (CaD_90_ and ContD_90_ respectively) were also significantly prolonged in the KCNH2 variant lines compared to the wildtype hiPSC-CMs (KCNH2^WT/WT^: 570±8ms (CaD_90_), 429±18ms (ContD_90_); KCNH2^PR/WT^: 620±10ms (CaD_90_), 492±23ms (ContD_90_); KCNH2^TL/WT^: 610±9ms (CaD_90_), 456±6ms (ContD_90_)).

Importantly, the APD_90_ and ContD_90_ of the KCNH2^PR/WT^ hiPSC-CMs were significantly prolonged compared to the KCNH2^TL/WT^ hiPSC-CMs. Although the CaD_90_ also appeared prolonged, this did not reach significance (*P*=0.05). This demonstrates that, under carefully controlled recording conditions, differences in I_Kr_ density are also reflected in the electrophysiological phenotype of the hiPSC-CMs, with the dominant negative-causing *KCNH2* mutation leading to a more pronounced increase in APD than the haploinsufficiency-causing *KCNH2* mutation. Furthermore, these differences are also reflected in both intracellular Ca^2+^ transients and contraction-relaxation kinetics, suggesting that these parameters are also differentially influenced by mild and severe LQT2-causing mutations.

### 3.4 KCNH2^PR/WT^ and KCNH2^TL/WT^ hiPSC-CMs exhibit differing sensitivities to E-4031

To determine if the electrophysiological differences observed between the three lines also led to differing responses to known arrhythmogenic compounds, we examined the response of the hiPSC-CMs to E-4031 (Figure 4). Figure 4A shows representative FP recordings in the presence of increasing concentrations of E-4031, with arrhythmic events such as EADs or fibrillations, detected in all three lines. As spontaneous beating ceased in some recordings when the cells were exposed to >300 nM E-4031 (Supplementary Figure S8), analysis of the effect of E-4031 on FPD prolongation was performed up to this concentration. The FPD of KCNH2^PR/WT^ and KCNH2^TL/WT^ hiPSC-CMs was significantly prolonged compared to KCNH2^WT/WT^ hiPSC-CMs at >1 nM E-4031 (Figure 4B). When FPD was normalised to baseline measurements, the change in FPD at 300 nM, was significantly different for the KCNH2^PR/WT^ hiPSC-CMs compared to other two lines (Figure 4C), indicating the KCNH2^PR/WT^ hiPSC-CMs were more sensitive to I_Kr_ blockade. Although the beat period between cell lines varied, it was unaffected for both the KCNH2^TL/WT^ and KCNH2^WT/WT^ hiPSC-CMs at <10 µM E-4031 (Supplementary Figure S9A). The cFPD was also examined, with differences between the KCNH2^PR/WT^ hiPSC-CMs and the other two lines still discernible (Supplementary Figure S9B-E).

**Figure 4.**
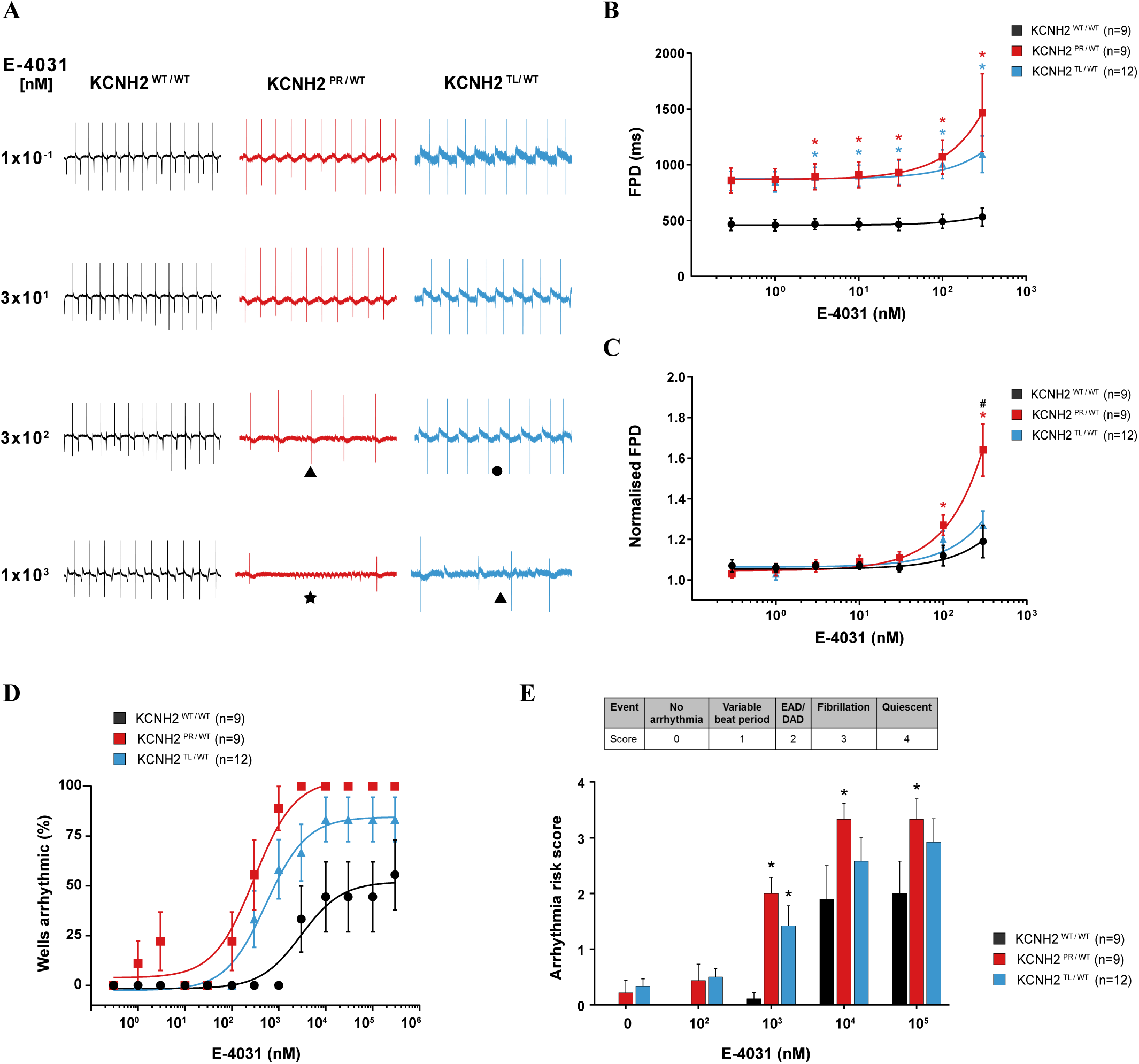
Effect of *I*_Kr_ blockade on FPD and arrhythmogenesis in KCNH2^WT/WT^, KCNH2^PR/WT^ and KCNH2^TL/WT^ hiPSC-CMs. **(A)** Representative MEA traces highlighting the difference in response upon accumulative addition of E-4031 between the indicated lines. Symbols indicate examples of the different types of arrhythmias detected: (•) variable beat period; (▴) abnormal repolarisations; (★) fibrillation. **(B, C)** FPD **(B)** and FPD normalised to baseline **(C)** of the indicated lines upon accumulative addition of E-4031. * indicates statistical significance to KCNH2^WT/WT^ (FPD: 3-300nM, *P*<0.05; normalised FPD: 100nM, *P*<0.05; 300nM, *P*<0.0001); # indicates statistical significance between KCNH2^PR/WT^ and KCNH2^TL/WT^ (*P*<0.0001); two-way ANOVA with Tukey’s multiple comparisons for *post hoc* test. **(D)** Scatter plot illustrating relationship between occurrence of arrhythmic events and concentration of E-4031 for the indicated lines. Curve fitting with nonlinear regression. **(E)** Arrhythmia risk scoring system and bar graph summarising the arrhythmia risk for each of the cell lines at different concentrations of E-4031. DAD, delayed afterdepolarisation; * indicates statistical significance to KCNH2^WT/WT^ (*P*<0.05; two-way ANOVA with Tukey’s multiple comparisons for *post hoc* test).

The second KCNH2^PR/WT^ and KCNH2^TL/WT^ clones also showed similar differences in sensitivity to E-4031 following FPD normalisation, with analysis performed to 100 nM E-4031 due to some KCNH2^PR/WT^ hiPSC-CMs becoming quiescent at 300 nM (Supplementary Figure S8; Figure S10A and B). Analysis of cFPD showed a similar trend although it was not significant (Supplementary Figure S10C, D), possibly due to overcompensation of the FPD-beat rate dependence for the KCNH2^TL/WT^ cl2 hiPSC-CMs (Supplementary Figure S10E-G). Overall, the multiple analyses we have performed with two separate clones for each mutation clearly demonstrate that the KCNH2-A561T mutation results in hiPSC-CMs that are more sensitive to I_Kr_ blockade than the KCNH2-N996I mutation.

To determine if these E-4031-induced differences in FPD between the lines also led to changes in the frequency of arrhythmia-like events, we examined the FP recordings for the occurrence of abnormal repolarisations, fibrillation, and quiescence. Persistent E-4031-induced arrhythmic events were first detected in the KCNH2^PR/WT^, KCNH2^TL/WT^ and KCNH2^WT/WT^ hiPSC-CMs at 100nM, 300nM and 1 µM respectively (Supplementary Figure S8). We also quantified the proportion of recordings that exhibited these arrhythmic responses with increasing concentrations of E-4031 (Figure 4A and D). Here too the KCNH2^PR/WT^ hiPSC-CMs were the most predisposed with 100% of recordings showing such events at ≥3 µM E-4031, followed by the KCNH2^TL/WT^ hiPSC-CMs with >80% of recordings becoming arrhythmic; while <55% of KCNH2^WT/WT^ hiPSC-CMs recordings showed such a response even at the highest E-4031 concentration (300µM). The E-4031 concentration that resulted in 50% of the maximal response was also significantly different between the lines (KCNH2^PR/WT^: 298 nM; KCNH2^TL/WT^: 536 nM; KCNH2^WT/WT^: 2.98 µM; *P*<0.01). This increased susceptibility to E-4031-induced arrhythmia-like events in the KCNH2^PR/WT^ hiPSC-CMs was also observed in the second set of KCNH2^PR/WT^ and KCNH2^TL/WT^ clones examined (Supplementary Figure S11A).

Finally, we investigated the possibility of developing a scoring system based on previously described methods ^27,28^ to estimate the arrhythmogenic risk to E-4031 for the different hiPSC-CM lines. We included variable beat period (score 1), which has previously been classified as a “mild” arrhythmia-type ^27^ as an additional category. Abnormal repolarisation was given a score of 2, while hiPSC-CMs that were fibrillating or became quiescent were scored 3 and 4 respectively. Both KCNH2^PR/WT^ and KCNH2^TL/WT^ hiPSC-CMs had higher arrhythmia risk scores compared to the KCNH2^WT/WT^ hiPSC-CMs at all concentrations of E-4031 analysed, with both lines significantly greater at 1 µM E-4031 and these differences remaining between the KCNH2^PR/WT^ and KCNH2^WT/WT^ hiPSC-CMs at higher concentrations (Figure 4E). Differences in the arrhythmia risk score were also observed between the second KCNH2^PR/WT^ and KCNH2^TL/WT^ clones (Supplementary Figure S11B). Taken together these results demonstrate a difference in susceptibility to arrhythmias between the variant lines and the KCNH2^WT/WT^ hiPSC-CMs, with KCNH2^PR/WT^ hiPSC-CMs more sensitive to E-4031 than the KCNH2^TL/WT^ hiPSC-CMs.

## 4. Discussion

Next-generation sequencing has had a profound impact on clinical genetic testing for LQTS, leading to the rapid identification of disease-causing variants in patients ^5^. However, interpreting the functional consequences of these variants is often inconclusive due to the variable expressivity and incomplete penetrance of these diseases ^4^, as well as the unexpectedly high level of background genetic variation observed in LQTS-susceptibility genes ^29^. This makes it challenging to not only accurately diagnose the patient, but also to determine appropriate clinical management. The ability to generate hiPSCs from patients, combined with advances in genome editing technologies, has demonstrated how such a platform can be used to determine the pathogenicity of variants of uncertain significance (VUS) ^30,31^, or the contribution of genetic modifiers to the disease phenotype ^32^. However, the extent to which hiPSCs can reflect intragenotype differences in disease risk such as that observed between LQT2 patients has not been fully explored ^33^. Here, we investigated this and demonstrate that genetically-matched hiPSC lines can model subtle differences in disease severity that are directly attributable to the *KCNH2* mutation.

Most *KCNH2* mutations in the cytoplasmic tail cause haploinsufficiency, with mutant subunits failing to co-assemble with wildtype subunits. This results in approximately half of the channels trafficking to the cell membrane and a *≤*50% reduction in I_Kr_ ^34^ In contrast, *KCNH2* mutations within the pore region typically have dominant-negative effects ^13^, frequently causing abnormal trafficking of the entire channel to the cell membrane, and thereby resulting in even fewer functional channels and a >50% I_Kr_ reduction ^34,35^. Consequently, LQT2 patients with missense mutations in the pore-loop region are at greater risk of an arrhythmia-related cardiac event ^9–12^. In the hiPSC-CMs we detected a <50% I_Kr_ decrease with the KCNH2-N996I mutation and a >70% reduction for the KCNH2-A561T mutation, indicating haploinsufficiency and dominant negative effects respectively, but also suggesting that tetrameric ion channels containing one KCNH2-A561T subunit can traffic to the cell membrane but have limited functionality. A previous study evaluating KCNH2-A561T in HEK293 cells was unable to detect any current ^36^, but here only the mutant KCNH2-A561T subunit was overexpressed which could explain why no channels were able to traffic to the membrane. Our findings also support other studies using hiPSCs that found pharmacological correction of trafficking defects may be a valid therapeutic strategy for *KCNH2* mutations located in the pore region ^18,37^.

We hypothesised that these differences in I_Kr_ would also be reflected in the electrophysiological phenotype of the hiPSC-CMs. Although all three platforms used to assess the electrophysiology of the lines demonstrated a clear prolongation of the APD and FPD for both KCNH2 variant lines compared with the KCNH2^WT/WT^ hiPSC-CMs, it was only with optical recordings that the APD was observed to be significantly more prolonged in KCNH2^PR/WT^ than KCNH2^TL/WT^ hiPSC-CMs. These discrepancies in sensitivity are likely due to differences in the setup and cellular configuration between the experimental approaches. Sparsely seeded hiPSC-CMs, such as those used for patch clamp recordings, show greater electrophysiological variability compared to measurements performed on confluent monolayers ^38^, which could confound the detection of subtle electrophysiological differences. Indeed, when confluent monolayers of hiPSC-CMs were measured using either MEAs or optically, the KCNH2^PR/WT^ lines had longer field- and action potentials compared to the KCNH2^TL/WT^ hiPSC-CMs that were significantly different when the cells were paced.

We also evaluated the cytosolic Ca^2+^ transients and contraction kinetics of all three lines, as these are also key parameters that can be altered in LQTS ^39–42^. As expected, both LQT2-causing mutations resulted in a significant prolongation in Ca^2+^ transients and contraction-relaxation duration when compared to KCNH2^WT/WT^ hiPSC-CMs. Interestingly, we also observed significant differences in contractility kinetics between the two KCNH2 variants. Contraction duration differences are known to exist between symptomatic and asymptomatic LQTS patients ^43^ and this has been proposed as an additional parameter to measure alongside QT interval for improving risk stratification in LQTS patients ^44^.

In LQT2 patients, sudden arousal is the most frequent trigger of an arrhythmic cardiac event ^45^, and patients with mutations in the pore-loop region have a significantly increased risk ^46^. This is also the case when the triggering factor is not arousal or exercise-related (e.g. fever, medication, sleep) ^46^. To mimic the effect of such triggers, we examined the behaviour of the hiPSC-CMs when treated with the I_Kr_ blocker, E-4031. We observed differing responses to the QT-prolonging drug between the three lines, with the KCNH2^PR/WT^ hiPSC-CMs exhibiting a larger response to I_Kr_ blockade than the KCNH2^TL/WT^ hiPSC-CMs, which exhibited similar changes in normalised FPD as KCNH2^WT/WT^ hiPSC-CMs. These differences in sensitivity to I_Kr_ block could be due to the dominant-negative effect of the KCNH2-PR mutation, additional gating kinetic defects in trafficked hERG channels that include the KCNH2-A561T subunit ^47^, or a combination of both. The results also differ from those recently reported by Yoshinaga and colleagues who observed a smaller change in FPD in response to I_Kr_ blockade in hiPSC-CMs derived from LQT2 patients than in control or mutation-corrected hiPSC-CMs ^48^. However, this discordance could be mutation-specific as they also observed an increased susceptibility to arrhythmias with the LQT2 hiPSC-CMs.

Indeed, in line with their differing FPD response to I_Kr_ blockade, the KCNH2^PR/WT^ hiPSC-CMs also exhibited an increased occurrence of arrhythmia events in the presence of E-4031. Similarly, a greater proportion of measurements from KCNH2^TL/WT^ hiPSC-CMs displayed arrhythmic activity compared to KCNH2^WT/WT^ hiPSC-CMs. Akin to systems being established to evaluate the arrhythmogenic risk of pharmacological compounds ^27^, a similar matrix could be developed to assess the risk of specific mutations in patients to different triggering conditions. As proof of concept, we determined the arrhythmia risk score for all three lines at various concentrations of E-4031, observing that the KCNH2^PR/WT^ hiPSC-CMs had a higher arrhythmogenic risk at 1 µM. It will be necessary to evaluate the accuracy and reliability of this scoring system with a larger panel of *KCNH2* mutations as well as for different triggers, but this study demonstrates the sensitivity of this *in vitro* model for detecting these intragenotype-phenotype differences. Such an approach could have important clinical implications, for example by identifying particular *KCNH2* mutations that predispose patients to increased arrhythmic risk and whom might benefit from more vigilant monitoring.

This study also highlights the benefit of introducing mutations into a well-characterised control hiPSC line. Indeed, this approach is the only option for identifying mutation-specific functional changes, as genetic background differences between patient-derived hiPSCs would likely mask these, even if the variant was corrected. This strategy also means that the mutations examined are not limited by the availability of patient material. While common variants present in the control hiPSC line might modify the disease phenotype, all mutations are evaluated on the same genetic background, thereby nullifying their effect. Furthermore, as similar responses were detected in both clonal lines generated for each mutation, the differences observed between the experimental groups are unlikely to be due to variants that have arisen spontaneously in culture or from CRISPR/Cas9-induced off-target effects.

In conclusion, we have established that hiPSCs can determine the underlying severity of individual *KCNH2* mutations, reflecting the mutation-location risk differences seen in LQT2 patients. This study demonstrates a new application in which hiPSC-CMs could contribute to improvements in the diagnosis, prognosis, and risk stratification of patients with congenital LQTS.

## Authors’ contributions

Conceptualization: KB and RD. Methodology: KB, LB, DM, CG, BM, MM and RD. Investigation: KB, LB, DM, AV and RD. Supervision: CM and RD. Formal analysis: KB, LB, DM, LS and RD. Software: BM and LT. Writing–Original draft: KB and RD. Writing–Review, Editing: KB, LB, DM, CG, BM, CM, LS and RD. Funding acquisition: RD

## Acknowledgements

We thank Maria Gomes Fernandes for assistance with immunofluorescence images, Jelle Goeman for statistical advice and Ana Krotenberg for technical assistance in performing the optical recordings. We also acknowledge Niels Geijsen for providing Cas9 protein.

