## Supplementary material for "Isogenic sets of hiPSC-CMs harboring *KCNH2* mutations capture location-related phenotypic differences"

**Short Title:** hiPSCs reveal LQT2 genotype-phenotype differences

**Karina O. Brandão[1], Lettine van den Brink[1], Duncan C. Miller[1], Catarina Grandela[1], Berend J. van Meer[1], Mervyn P.H. Mol[1], Leon G.J. Tertoolen[1], Christine L. Mummery[1], Luca Sala[2], Arie O. Verkerk[3, 4], Richard P. Davis[1*]**

1. Department of Anatomy and Embryology, Leiden University Medical Center, 2300 RC Leiden, The Netherlands;
2. Istituto Auxologico Italiano, IRCCS, Laboratory of Cardiovascular Genetics, 20095 Milan, Italy;
3. Department of Medical Biology, Amsterdam UMC, University of Amsterdam, 1105 AZ Amsterdam, The Netherlands;
4. Department of Experimental Cardiology, Amsterdam UMC, University of Amsterdam, 1105 AZ Amsterdam, The Netherlands

**Supplementary material contents:**

**11 Supplementary Figures + Legends**

**Supplementary Methods**

**Supplementary References**

**5 Supplementary Tables**

#### Supplementary Figures and Legends

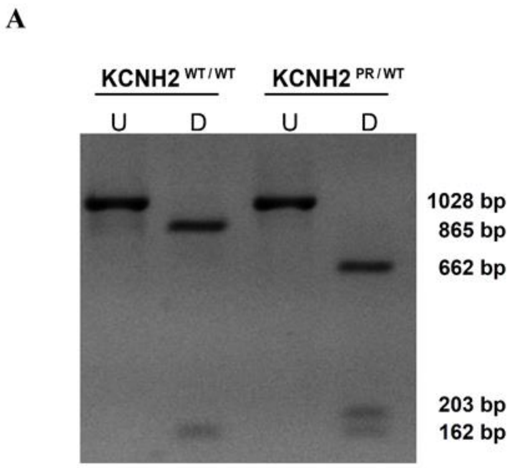

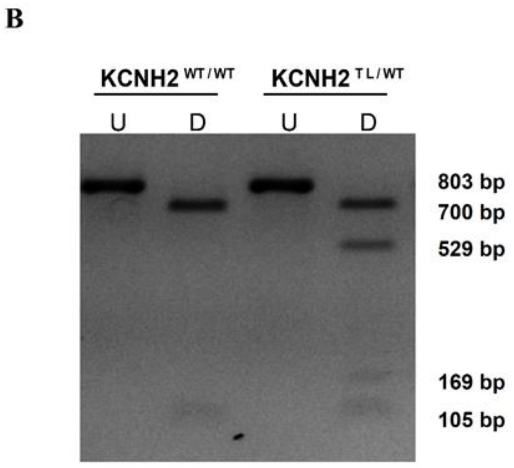

**Supplementary Figure 1: Restriction Fragment Length Polymorphism assay to detect targeted clones.** PCR products amplified from all lines were subjected to restriction digest with enzymes that recognize the site modified by CRISPR/Cas9, resulting in fragments of different sizes (indicated on the right of each image) between wild type and mutated lines. **(A)** KCNH2$^{WT/WT}$ and KCNH2$^{PR/WT}$ hiPSCs digested with *HaeII*. **(B)** KCNH2$^{WT/WT}$ and KCNH2$^{TL/WT}$ with *BccI* enzyme. U – undigested; D – digested; bp – base pairs

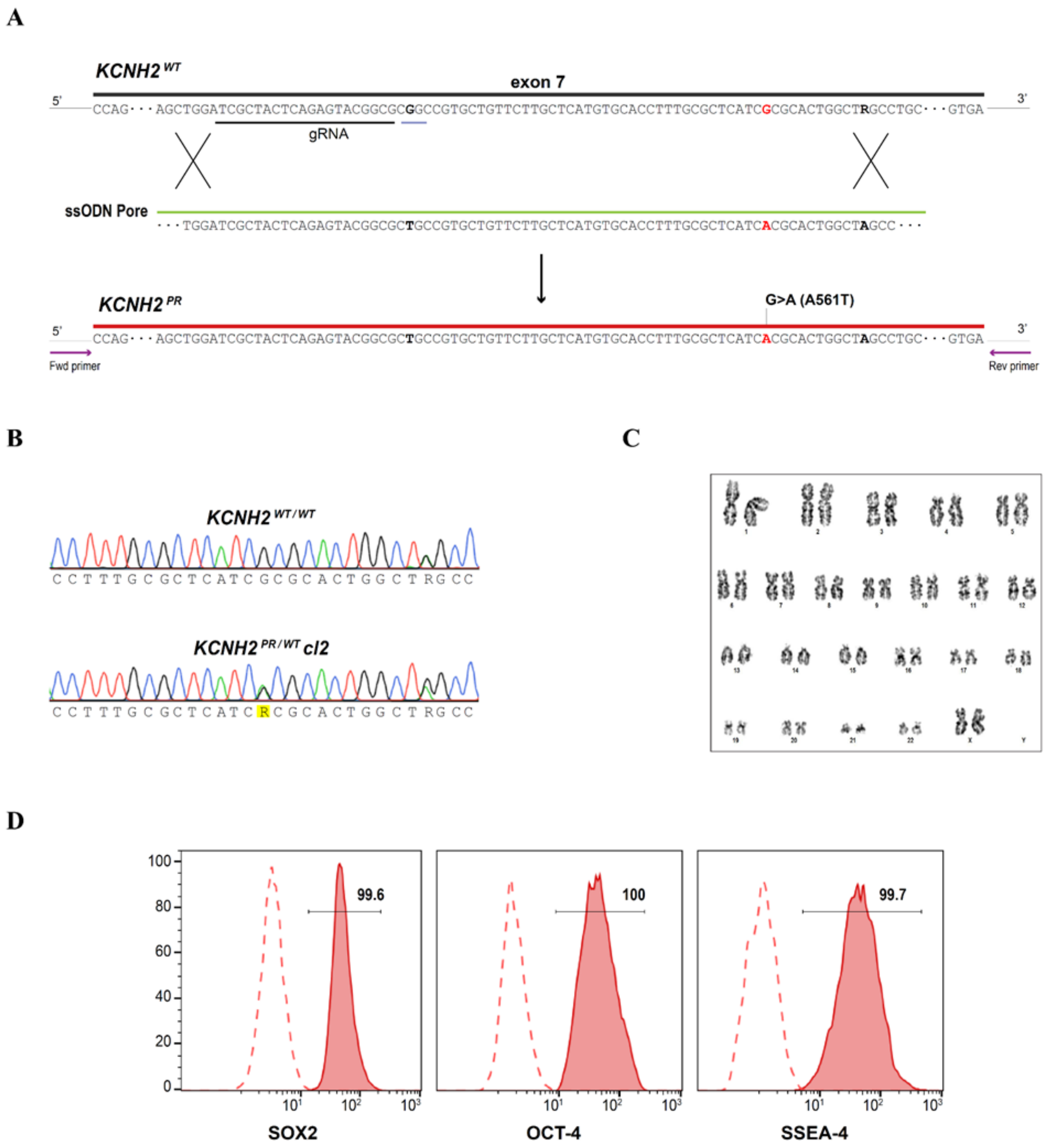

**Supplementary Figure 2: Generation of the KCNH2$^{PR/WT}$ cl2 hiPSC line. (A)** Schematic outlining the gene editing strategy to introduce the A561T mutation (*KCNH2$^{PR}$*) by homologous recombination into a *KCNH2* wildtype (*KCNH2$^{WT}$*) sequence. A gRNA (black underline) was designed ~40 nucleotides 5' of the mutation. Red nucleotide indicates nucleotide being mutated. Nucleotide in bold within the PAM sequence (grey underline) was modified to prevent Cas9 cutting the mutated sequence. Additional bold nucleotide corresponds to a heterozygous synonymous SNP (rs1805121). Primers (purple arrows) amplified ~1 kb surrounding the modified sequence for screening. **(B)** Sequence analysis of the PCR-amplified DNA showing heterozygous introduction of the c.G1681A (A561T) mutation (highlighted R) in the KCNH2$^{PR/WT}$ cl2 hiPSC line. **(C)** G-band karyogram indicating no chromosomal aberrations in the KCNH2$^{PR/WT}$ cl2 hiPSC line. **(D)** Flow cytometry analysis of the pluripotency-associated markers SOX2, OCT4 and SSEA4. Values in the histograms show the percentage of hiPSCs positive for the indicated marker. Dotted lines represent a negative control population.

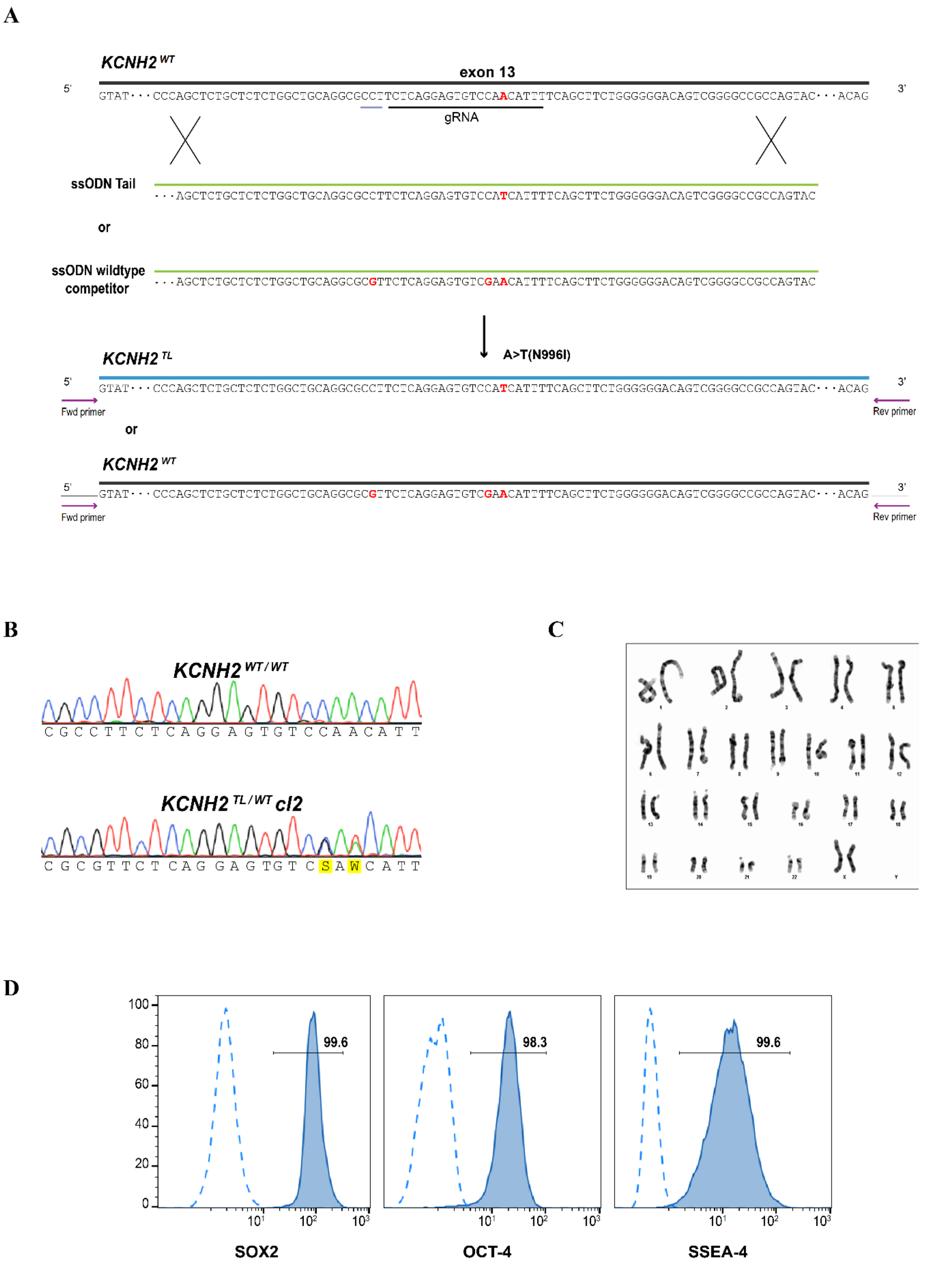

**Supplementary Figure 3: Generation of the KCNH2$^{TL/WT}$ cl2 hiPSC line. (A)** Schematic outlining the gene editing strategy to introduce the N996I mutation (*KCNH2$^{TL}$*) by homologous recombination into a *KCNH2* wildtype (*KCNH2$^{WT}$*) sequence. The gRNA (black underline) and Cas9 RNP were co-transfected with two ssODN sequences. The *ssODN Tail* included the c.A2987T (N996I) mutation (red), while the *ssODN wildtype competitor* included silent nucleotide mutations (red) that did not alter any amino acids but assisted with RFLP screening and prevented Cas9 from cutting the modified sequence due to a mutation in the PAM sequence (grey underline). By co-transfecting two ssODN templates, we hypothesised this would increase the frequency of generating a heterozygous KCNH2$^{TL/WT}$ hiPSC line. Primers (purple arrows) amplified ~1 kb surrounding the modified sequence for screening. **(B)** Sequence analysis of the PCR-amplified genomic DNA showing heterozygous introduction of the c.A2987T (N996I) mutation (highlighted W), as well as the RFLP screening silent mutation (highlighted S), in the KCNH2$^{TL/WT}$ cl2 hiPSC line. **(C)** G-band karyogram indicating no chromosomal aberrations in the KCNH2$^{TL/WT}$ cl2 hiPSC line. **(D)** Flow cytometry analysis of the pluripotency-associated markers SOX2, OCT4 and SSEA4. Values in the histograms show the percentage of hiPSCs positive for the indicated marker. Dotted lines represent a negative control population.

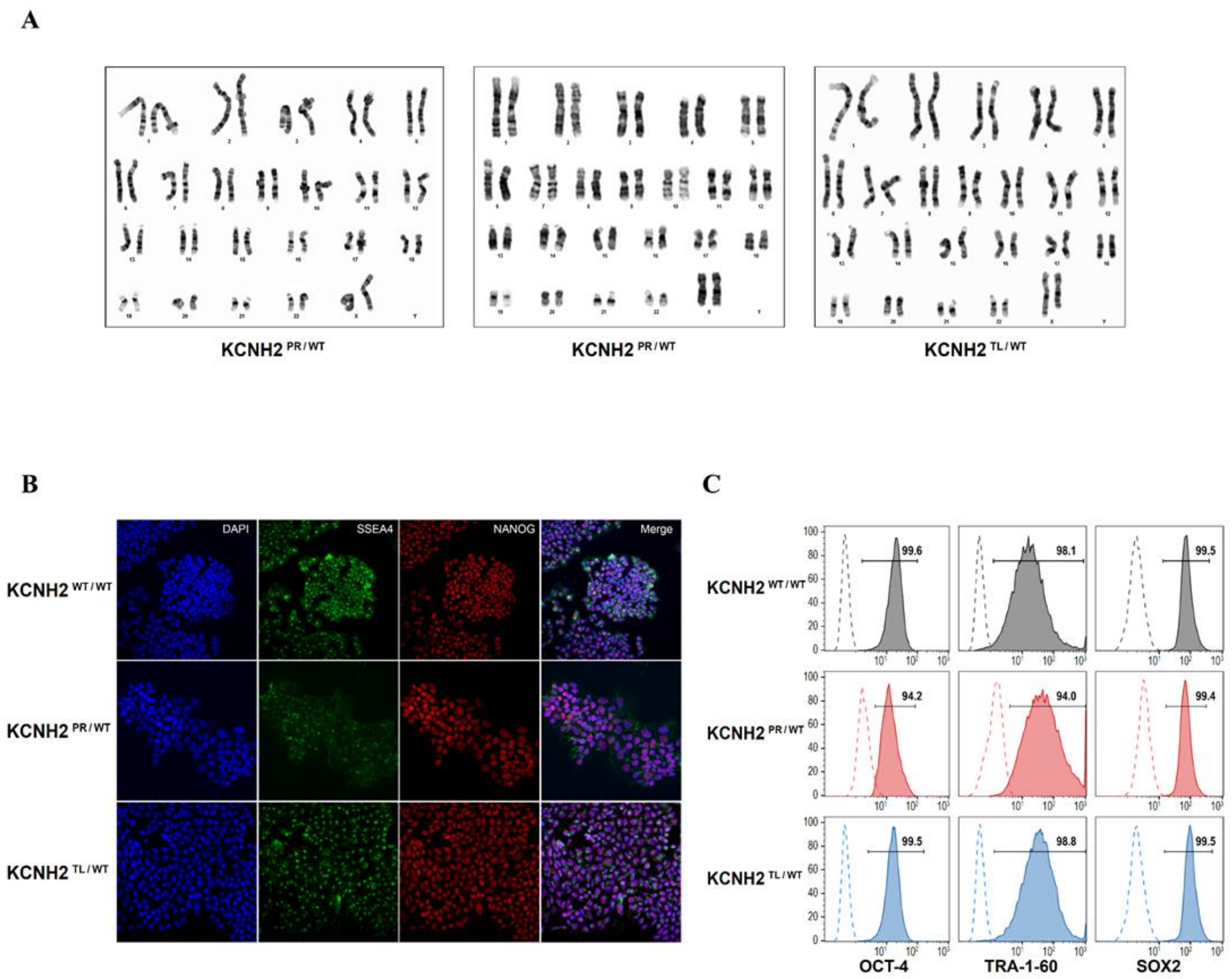

**Supplementary Figure 4: Characterisation of the set of isogenic *KCNH2* hiPSC lines. (A)** G-band karyograms indicating a normal euploid karyotype for KCNH2$^{WT/WT}$, KCNH2$^{PR/WT}$ and KCNH2$^{TL/WT}$ hiPSCs. **(B)** Immunofluorescence and **(C)** flow cytometry analysis of the pluripotency-associated markers, NANOG, OCT4, SOX2, TRA-1-60 and SSEA4 in the indicated hiPSC lines. Nuclei were stained with DAPI **(B)**, while dotted lines represent a negative control population **(C)**. Values in the histograms show the percentage of hiPSCs positive for the indicated marker.

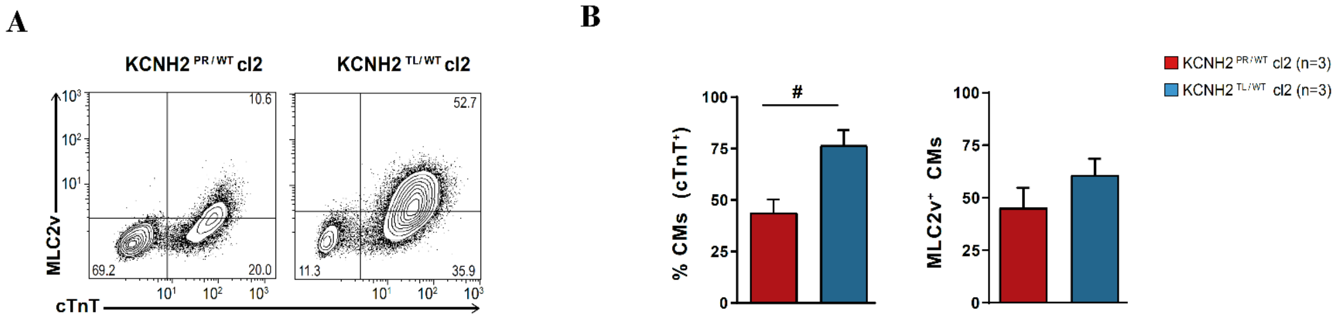

**Supplementary Figure 5: Characterisation of KCNH2$^{PR/WT}$ cl2 and KCNH2$^{TL/WT}$ cl2 hiPSC-CMs.** **(A)** Representative flow cytometry plots of hiPSC-CMs in the indicated lines for expression of cTnT and MLC2v, 7 days after replating. Values inside the plots are the percentage of cells within the gated region. **(B)** Overall cardiac differentiation efficiency of the two hiPSC lines. While the percentage of hiPSC-CMs (cTnT$^{+}$) differentiated from the KCNH2$^{TL/WT}$ cl2 hiPSCs was similar (77%) to the other hiPSC lines, the percentage of CMs obtained from the KCNH2$^{PR/WT}$ cl2 hiPSCs was less ($P$=0.02; left bar graph). However, the proportion of ventricular-like cardiomyocytes in the hiPSC-CM population was not significantly different between the two clones and was also similar to the other hiPSC lines (right bar graph).

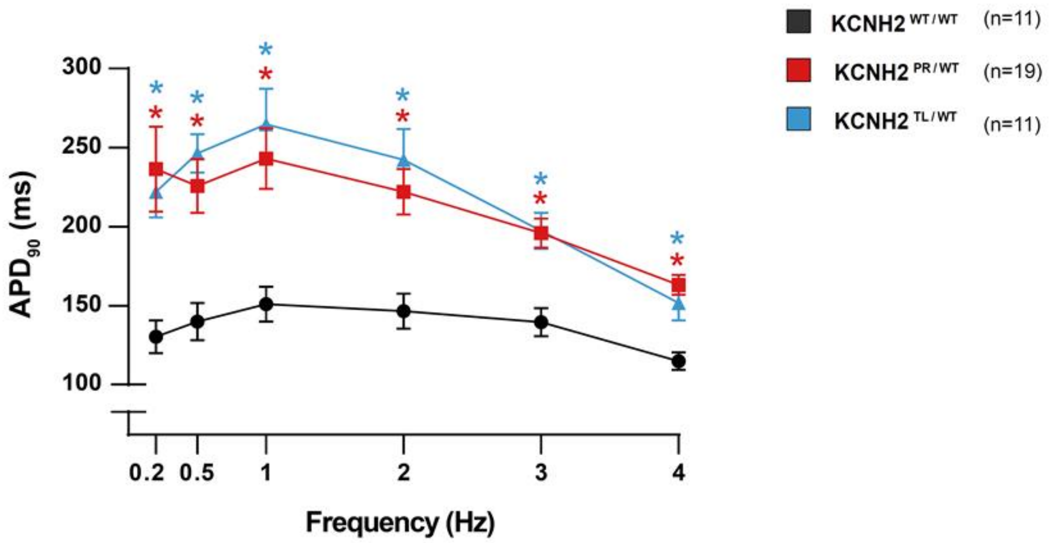

**Supplementary Figure 6:** Average AP duration at 90% repolarisation ($APD_{90}$) values for KCNH2$^{PR/WT}$ and KCNH2$^{TL/WT}$ hiPSC-CMs show that the lines have a significantly prolonged $APD_{90}$ at all pacing frequencies between 0.2–4 Hz compared to KCNH2$^{WT/WT}$ hiPSC-CMs. * indicates statistical significance to KCNH2$^{WT/WT}$ ($P$<0.01).

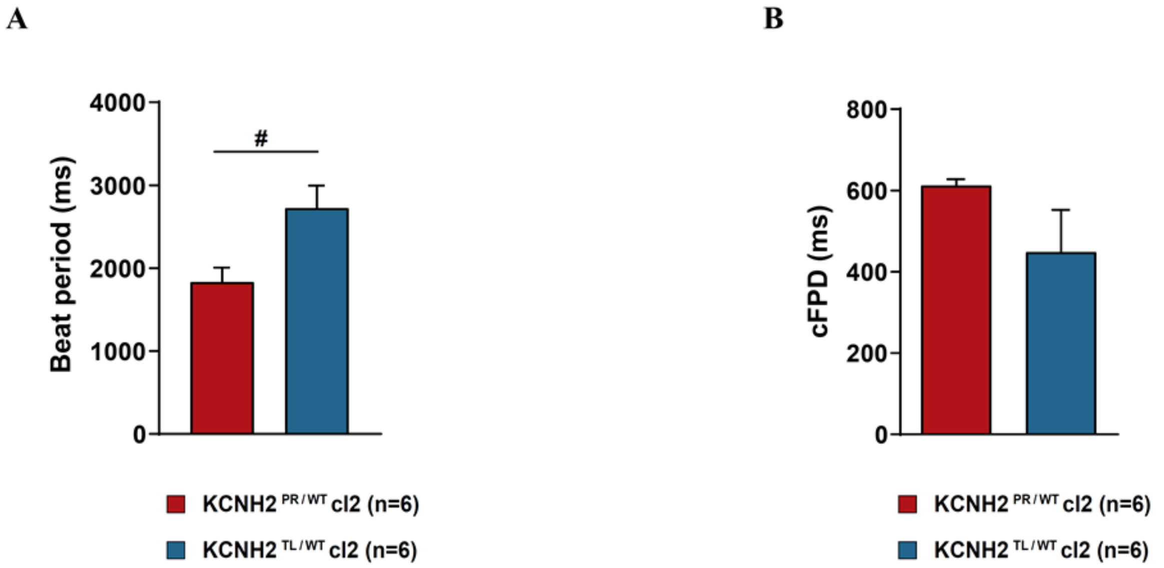

**Supplementary Figure 7: Electrophysiological characterisation of KCNH2$^{PR/WT}$ cl2 and KCNH2$^{TL/WT}$ cl2 hiPSC-CMs using multielectrode arrays.** Average values for beat period interval **(A)** and the cFPD **(B)** for the indicated cell lines. # indicates statistical significance between mutated lines ($P$<0.05).

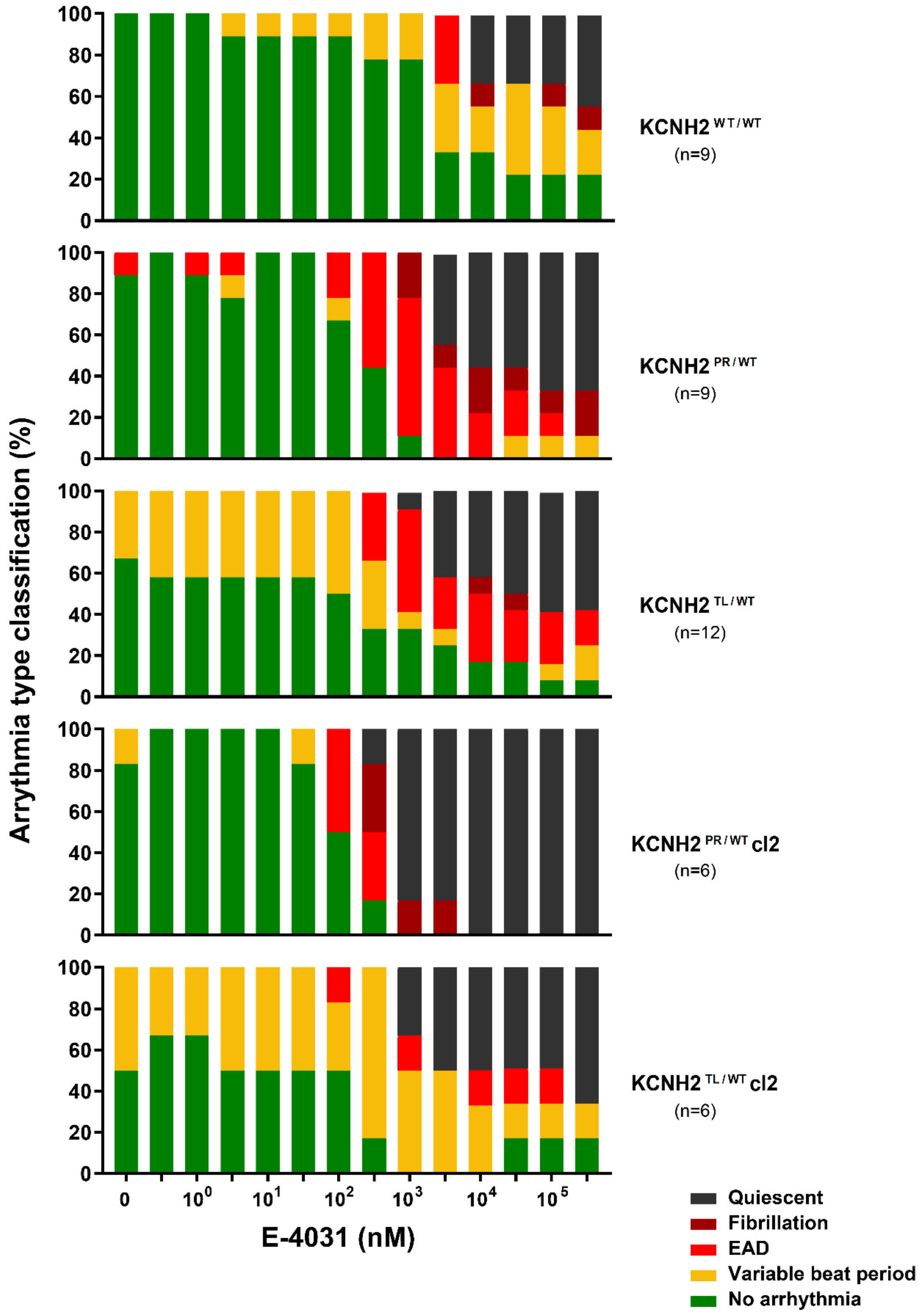

**Supplementary Figure 8:** Quantification of the arrhythmia subtypes detected with increasing concentrations of E-4031 for the indicated lines.

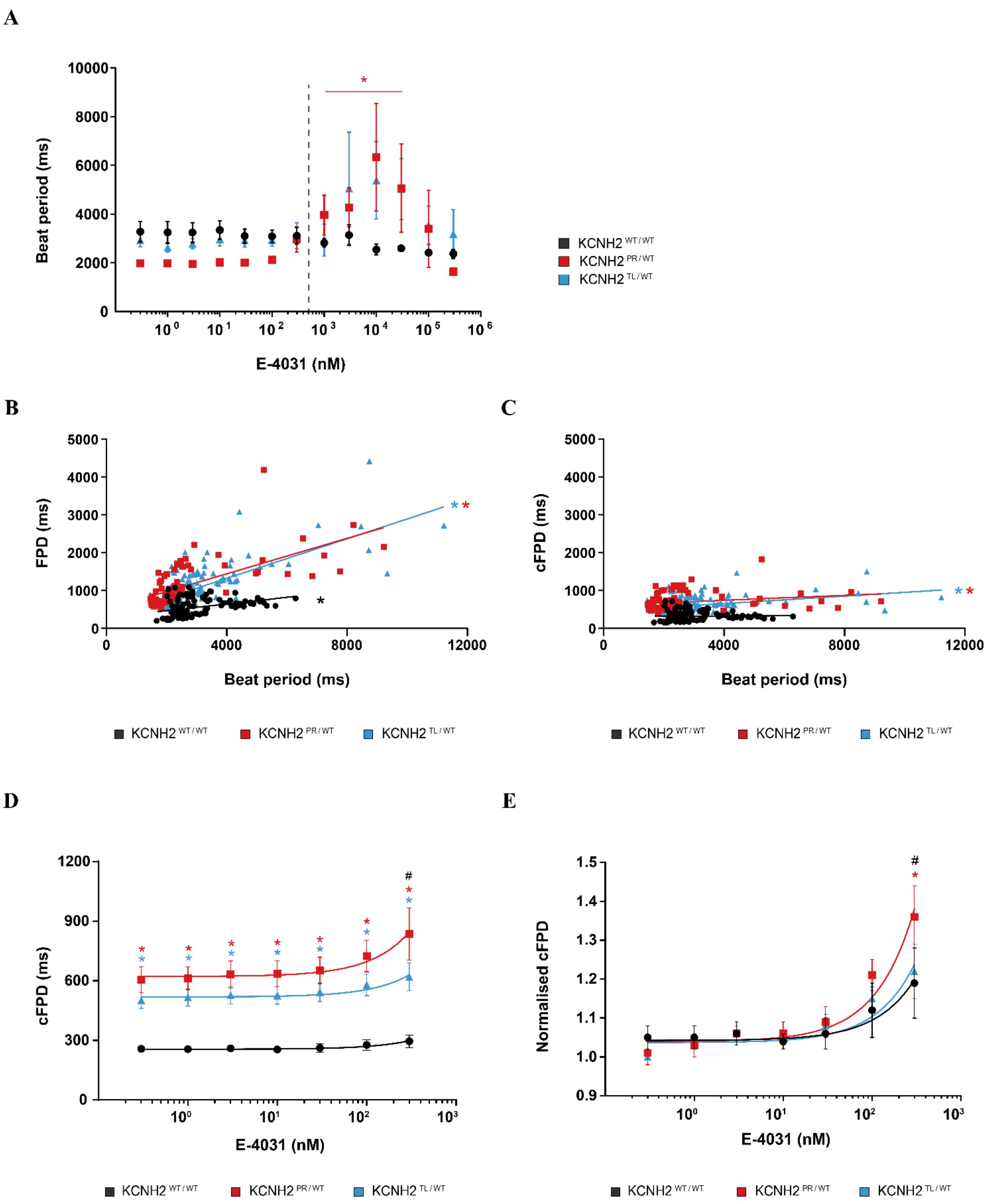
A
B
C
D
E
Beat period (ms)
E-4031 (nM)
FPD (ms)
cFPD (ms)
Normalised cFPD
KCNH2 WT / WT
KCNH2 PR / WT
KCNH2 TL / WT
*
#

**Supplementary Figure 9: Analysis of the effect of E-4031 on beat period and FPD of the KCNH2$^{WT/WT}$, KCNH2$^{PR/WT}$ and KCNH2$^{TL/WT}$ hiPSC-CMs. (A)** Average beat period for the indicated lines upon increasing concentrations of E-4031 (0.3 nM - 300 µM). The dotted line demarcates the maximum concentration of E-4031 (300 nM) included in the analysis of the effect of this compound on FPD in Figure 4B and 4C. * indicates statistical significance of a cell line to its respective baseline ($P$<0.05). **(B & C)** Major-axis regression analysis on the relationship between beat period and either FPD **(B)** or cFPD **(C)**, for the indicated lines upon increasing concentrations of E-4031 (0.3 nM - 300 µM). * indicates linear coefficients significantly different from 0 ($P$<0.0001). **(D & E)** cFPD **(D)** and cFPD normalised to baseline **(E)** of the indicated lines upon accumulative addition of E-4031 (0.3-300 nM). * indicates statistical significance to KCNH2$^{WT/WT}$ (cFPD: 0.3-300 nM, $P$<0.02; normalised cFPD: 300 nM, $P$<0.05); # indicates statistical significance between KCNH2$^{PR/WT}$ and KCNH2$^{TL/WT}$ hiPSC-CMs ($P$<0.05); KCNH2$^{WT/WT}$ and KCNH2$^{PR/WT}$, n=9; KCNH2$^{TL/WT}$, n=12.

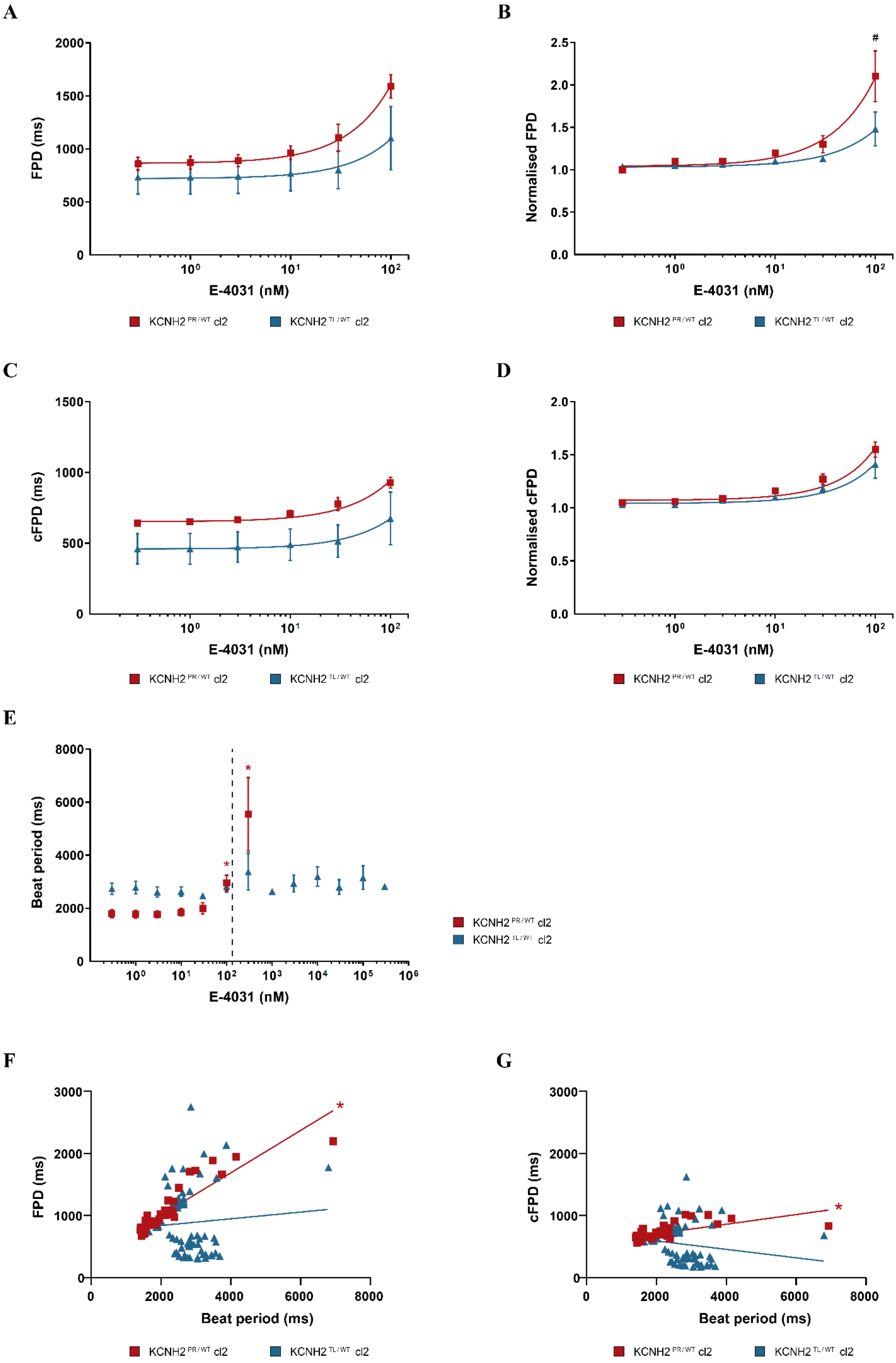
A
B
C
D
E
F
G
FPD (ms)
Normalised FPD
cFPD (ms)
Normalised cFPD
Beat period (ms)
E-4031 (nM)
#
*
KCNH2 PR/WT cl2
KCNH2 TL/WT cl2

**Supplementary Figure 10: Effect of $I_{Kr}$ blockade on (c)FPD in hiPSC-CMs from a second $KCNH2^{PR/WT}$ and $KCNH2^{TL/WT}$ clone. (A-D)** FPD **(A)**, FPD normalised to baseline **(B)**, cFPD **(C)** and cFPD normalised to baseline **(D)** of the indicated lines upon accumulative addition of E-4031. # indicates statistical significance between mutated lines ($P<0.05$); n=6 for $KCNH2^{PR/WT}$ cl2 and $KCNH2^{TL/WT}$ cl2. **(E)** Average beat period for the indicated lines upon addition of increasing amounts of E-4031 (0.3 nM - 300 µM). The dotted line demarcates the maximum concentration of E-4031 (100 nM) included in the analysis shown in **(A-D).** * indicates statistical significance for indicated line to its respective baseline ($P<0.01$). All $KCNH2^{PR/WT}$ samples became quiescent at ≥10 µM of E-4031. **(F & G)** Major-axis regression analysis on the relationship between beat period and either FPD **(F)**, or cFPD **(G)**, for the indicated lines upon addition of increasing amounts of E-4031 (0.3 nM - 300 µM). * indicates linear coefficients significantly different from 0 ($P<0.0001$).

A

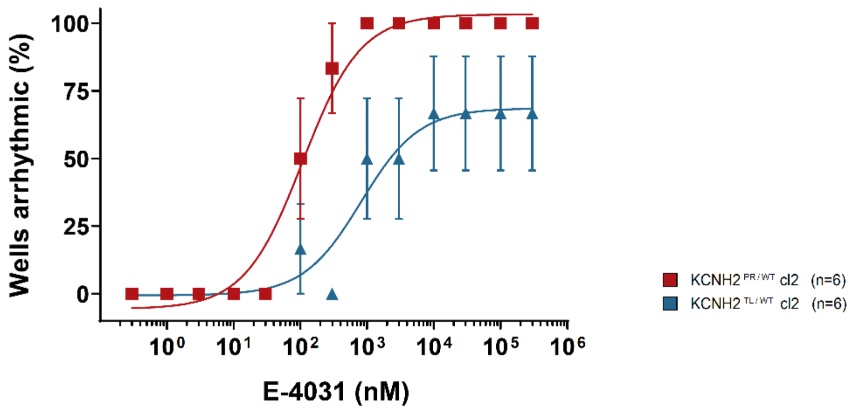

B

| Event | No arrhythmia | Variable beat period | EAD/ DAD | Fibrillation | Quiescent |
|---|---|---|---|---|---|
| Score | 0 | 1 | 2 | 3 | 4 |

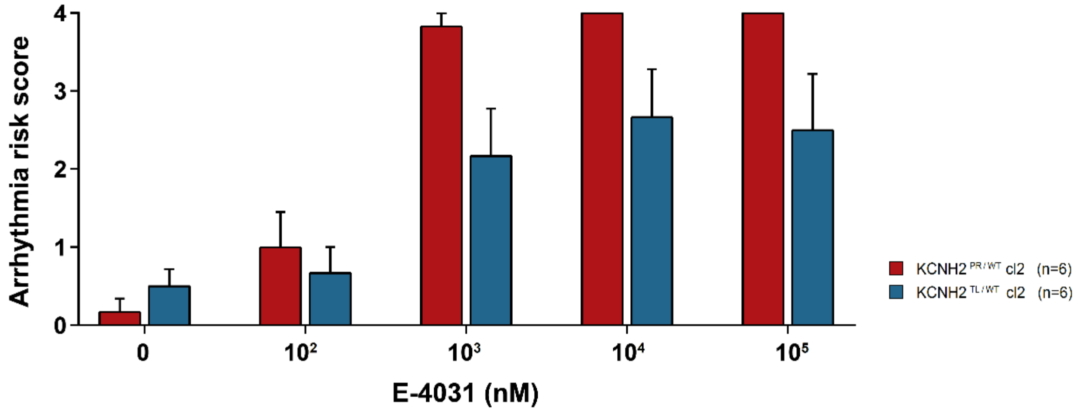

**Supplementary Figure 11: Evaluation of arrhythmic events in KCNH2$^{PR/WT}$ cl2 and KCNH2$^{TL/WT}$ cl2 hiPSC-CMs in response to increasing concentrations of E-4031. (A)** Scatter plot illustrating relationship between occurrence of arrhythmic events and concentration of the $I_{Kr}$ blocker for the indicated lines. For each cell line, the percentage of wells displaying an arrhythmic episode was plotted against the corresponding drug concentration. Curve fitting with nonlinear regression. **(B)** Arrhythmia risk scoring system and bar graph summarising the arrhythmia risk for each of the cell lines at different concentrations of E-4031.

### Supplementary Methods

##### Culture of human induced pluripotent stem cells (hiPSCs)

For maintenance of the hiPSC lines, Essential 8™ Medium (Gibco) and Vitronectin (VTN-N; Gibco)-coated plates were used. For passaging, the cells were dissociated with TrypLE Select Enzyme (Gibco) and re-plated in Essential 8™ Medium containing RevitaCell™ Supplement (1:200 dilution; Gibco). For gene editing experiments, the hiPSCs were maintained either on irradiated mouse embryonic fibroblasts in human embryonic stem cell medium as previously described[1] or in StemFlex™ Medium (Gibco) according to the manufacturer's instructions.

##### Differentiation and culture of hiPSC-derived cardiomyocytes (hiPSC-CMs)

One day prior to differentiation, hiPSCs were seeded at 3.75 x $10^4$/$cm^2$ into Matrigel® Matrix (BD)-coated wells in Essential 8™ Medium containing RevitaCell™ Supplement. The hiPSCs were differentiated into cardiomyocytes using the Pluricyte Cardiomyocyte Differentiation Kit (NCardia) according to the manufacturer's instructions, with cells maintained in Medium C until day 19-21 of differentiation. The hiPSC-CMs were dissociated with 5x TrypLE Select Enzyme for 10 min at 37°C and cryopreserved. For replating the hiPSC-CMs, the frozen cells were thawed at 37°C and transferred to a conical tube. Immediately thereafter, 1 ml of BPEL medium[2] was added dropwise (1 drop every 5 s), followed by ~4.7 ml BPEL (1 drop every 2 s). Cells were pelleted at 250*g* for 3 min, resuspended in Medium C and plated as required.

##### Exome sequencing

Genomic DNA (gDNA) from $KCNH2^{WT/WT}$ hiPSCs was extracted using the Blood and Cultured Cells DNA Mini Kit (Qiagen), according to manufacturer's instructions. Whole exome sequencing was performed by BGI (Hong Kong) using a HiSeq X platform (Illumina), with the library constructed from 6 µg gDNA using the SureSelect Human All Exon V5 +UTR enrichment kit (Agilent). Bioinformatics analysis (Mendel disease) was performed by BGI. Exome sequencing coverage and variant calls were manually curated for 107 genes known to be linked to inherited arrhythmia syndromes or cardiomyopathies[3].

**Generation of the guide RNAs (gRNAs)**

Candidate gRNAs were identified around the intended mutation site using the bioinformatic website http://crispor.tefor.net, and gRNAs with higher specificity were favoured to minimise potential off-target sites (Supplementary Table S1). The gRNAs were either synthesised as chimeric single gRNA (sgRNA) by *in vitro* transcription or purchased as crRNA and complexed with tracrRNA (both from IDT) prior to transfection.

For *in vitro* transcription, primers containing the T7 promoter, crRNA and a sequence homologous to the tracrRNA component present in the vector pSp-Cas9(BB)-2A-puro_v2 (Addgene #62988) were designed along with a primer complementary to the 3' end of the tracrRNA (Supplementary Table S2). A 110bp amplicon was generated by PCR and purified using the QiaQuick PCR purification kit (Qiagen). The sgRNAs were transcribed using the HiScribe T7 kit (New England Biolabs). Following purification using the NucleoSpin RNA Clean-up XS kit (Macherey-Nagel), the integrity of the sgRNAs was confirmed by electrophoresis using a 10% TBE-urea precast gel (Bio-Rad).

The crRNAs and tracrRNA were resuspended in Nuclease-Free Duplex Buffer (IDT) to a final concentration of 100 µM. For complexing the RNAs, a 20 µM reaction was prepared by mixing 1 µl of each RNA with 3 µl of nuclease-free duplex buffer, incubating at 95˚C for 5 min, and leaving to cool to room temperature for at least 2 h before transfection.

**Design of the single stranded oligonucleotides (ssODNs)**

Approximately 125 bp asymmetric oligonucleotides were designed to contain the desired mutations and to have at least 36 nucleotides homologous to the genomic sequence 3' of the Cas9-induced double strand break (Supplementary Table S1). The ssODNs were purchased as Ultramer DNA oligos (IDT) with standard desalting purification and resuspended in TE buffer to a final concentration of 100 µM.

**CRISPR/Cas9 and ssODN transfection into hiPSCs**

The gRNA, Cas9 protein (IDT or kindly provided by Niels Giessen[4]) and ssODN were transfected into the hiPSCs by either electroporation or lipofection. For electroporation, 1 µg Cas9 and 240 ng gRNA were mixed in a 0.5 ml sterile Protein LoBind tube (Eppendorf) and incubated at 25°C for 10 min to form the Cas9 ribonucleotide protein (RNP) complex. The hiPSCs were harvested and 1 x $10^5$ cells mixed with the Cas9 RNP and 40 pmol ssODN in a 10 µl total reaction. Electroporation was performed using the Neon Transfection System (ThermoFisher), and the electroporated cells transferred immediately into a Laminin-521 coated well containing 500 µl StemFlex™ Medium with RevitaCell™ Supplement (1:100). Approximately 3 days later, the cells were harvested and expanded for subcloning.

For lipofection, MEF-maintained hiPSCs were transfected when 60–75% confluent. On the day of transfection, medium was refreshed 1 h before the procedure. Transfection reactions were prepared in a 0.5 ml sterile Protein LoBind tube by mixing 1.5 µg Cas9 with 380 ng of sgRNA, followed by the addition of Opti-MEM™ I Reduced Serum Medium (Gibco) to a final volume of 25 µl, and incubated as described above to form the Cas9 RNP complex. For co-delivery of the donor template, 4 pmol of ssODN was diluted in Opti-MEM™ I Reduced Serum Medium to a final volume of 25 µl. A lipofectamine mixture was prepared by diluting Lipofectamine® 2000 Transfection Reagent (Invitrogen) in Opti-MEM™ I Reduced Serum Medium (1:10). The Cas9 RNP, and ssODN solution were then combined with the diluted lipofectamine to a final volume of 100 µl and incubated at room temperature for 10 min before added to the hiPSCs. Approximately 6 h after transfection the media was refreshed, and at 72 h, the cells harvested and expanded for subcloning.

**Targeting strategy to generate the KCNH2$^{PR/WT}$ hiPSC lines**

Using the electroporation method, the KCNH2$^{PR/WT}$ hiPSC lines were created by introducing a heterozygous point mutation to substitute alanine to threonine at position 561 of KCNH2. A gRNA site 45 nucleotides 5' of the mutation site was targeted with a crRNA-tracrRNA complex, and the ssODN (ssODN_Pore) was designed to have a silent mutation within the PAM sequence to aide both colony screening and to prevent re-cutting of the modified locus by Cas9. For screening the resulting colonies, approximately 1 kB surrounding the target site was amplified by PCR and digested with *HaeII* to identify putative mono-allelic targeted clones. These were subsequently confirmed to be heterozygous for the mutation by Sanger sequencing.

**Targeting strategy to generate the KCNH2$^{TL/WT}$ hiPSC lines**

The KCNH2$^{TL/WT}$ hiPSC line was generated by introducing a heterozygous point mutation to substitute asparagine to isoleucine at position 996 of KCNH2. The target sequence for the sgRNA included the site to mutate, and the ssODN (ssODN Tail) was designed to include the desired nucleotide modification. For the KCNH2$^{TL/WT}$ cl2 hiPSC line a different sgRNA also covering the mutation site was transfected, along with both the ssODN Tail as well as a ssODN lacking the mutation but including two silent mutations. This additional ssODN functioned as competitor homology template to improve the frequency of obtaining a heterozygous clone. For screening the resulting colonies, approximately 1 kB surrounding the target site was amplified by PCR and digested with *BccI* to identify putative mono-allelic targeted clones. These were subsequently confirmed to be heterozygous for the mutation by Sanger sequencing.

**Subcloning and PCR screening of the hiPSCs**

Transfected hiPSCs were clonally isolated by single-cell deposition using a flow cytometer. Briefly, the cells were harvested using TrypLE™ Select Enzyme and filtered to remove cell clumps. If the transfected hiPSCs had been cultured on MEFs, prior to subcloning, the cells were stained with anti-MEF antibody (PE-conjugated; Miltenyi Biotec) to exclude these cells. All lines were stained with 4’,6 Diamidino-2-Phenylindole (DAPI, Invitrogen) to exclude dead cells. A single hiPSC was deposited directly into each well of a 96-well plate in the same culture conditions as what the transfected hiPSCs had been maintained in. To assist with clonal recovery, the culture media also contained RevitaCell™ Supplement (1:100). Medium was initially changed three days after deposition and cells were maintained for ~2 weeks with medium changes every 3-4 days. Wells containing hiPSC colonies were dissociated with StemPro™ Accutase™ Cell Dissociation Reagent (Gibco) and duplicated across two 96-well plates. DNA for PCR screening and restriction fragment length polymorphism (RFLP) analysis was isolated from the cells in one plate with QuickExtract™ DNA Extraction Solution (Lucigen), while the cells in the replicate plate were cryopreserved.

**Off-target analysis**

Potential genomic off-targets for each gRNA were identified using http://crispor.tefor.net. The top 5 candidates with up to 4 base mismatches, were evaluated for the presence of Cas9-induced mutations (Supplementary Table S5). The region around the off-target site was PCR amplified from both the targeted and KCNH2$^{WT/WT}$ hiPSC lines. The PCR amplicons were treated with Exonuclease I and Shrimp Alkaline Phosphatase (both New England Biolabs) and sequenced by Sanger sequencing to confirm the absence of off-target modifications.

**Karyotyping**

The genetically-modified hiPSC lines were karyotyped by G-banding. Chromosome spreads and analyses were performed by Cell Guidance Systems (UK) or by the Laboratory for Diagnostic Genome Analysis (Leiden University Medical Center). For each cell line, 20 metaphase spreads were examined with samples of sufficient quality to detect numerical and large structural abnormalities.

**Allele-specific expression of *KCNH2***

Total RNA was isolated from hiPSC-CMs using the NucleoSpin® RNA Kit (Macherey-Nagel) according to manufacturer's instructions, with DNAse treatment performed using the DNA-free™ DNA Removal Kit (Ambion). RNA was reverse transcribed into cDNA using the iScript™ cDNA Synthesis Kit (Bio-Rad). Using primer pairs flanking each mutation (Supplementary Table S2), PCR products generated from the *KCNH2* transcripts were cloned into the pMiniT™ 2.0 vector using the NEB® PCR Cloning Kit (New England Biolabs). For each *KCNH2* variant line at least 29 clones containing the transcript underwent Sanger sequencing.

**Flow cytometric analysis**

A single cell suspension of hiPSCs or hiPSC-CMs was obtained by dissociating the cells with TrypLE™ Select Enzyme and filtering the cell suspension. Cells were fixed and permeabilised using the Fix & Perm Cell Permeabilization Kit (Invitrogen) according manufacturer's instructions. For the hiPSC-CMs, the cells were first incubated with a Viobility™ 405/520 fixable dye (Miltenyi Biotec) prior to fixation for subsequent exclusion of dead cells. The hiPSCs were incubated with the conjugated antibodies OCT4-BV421 (1:25, BD #565644), Sox2-A488 (1:200, eBioscience, #53-9811-80), Tra-1-60-PE (1:20, Miltenyi Biotec, #130-100-347) and SSEA4-PE-vio770 (1:100, Miltenyi Biotec, #130-105-082), while the hiPSC-CMs were incubated with cTnT-Vioblue (1:11, Miltenyi Biotec, #130-106-686) and MLC2v-PE (1:11, Miltenyi Biotec, #130-106-183). All antibodies were diluted in permeabilization medium (medium B; Invitrogen). Samples were measured using a MACSQuant VYB flow cytometer (Miltenyi Biotec), and data analysed using FlowJo software (FlowJo).

**Immunofluorescence analysis**

For hiPSCs, cells were fixed in 2% paraformaldehyde for 30 min, permeabilized with phosphate buffer saline (PBS)/0.1% Triton X-100 (Sigma-Aldrich) and blocked with 1% bovine serum albumin (BSA, Sigma-Aldrich) and 0.05% Tween (Merck) in PBS. Samples were incubated overnight at 4°C with antibodies specific for NANOG (1:200, R&D, #AF1997) and SSEA4 (1:200, Santa Cruz Biotechnology, #SC59368). These primary antibodies were detected with Alexa Fluor 555- (1:500, ThermoFisher, #A21432) and Alexa Fluor 488- (1:500, Life Technologies, #A-21202) conjugated antibodies, respectively. Nuclei were visualised with

DAPI (0.3 µM) and images captured using an EVOS FL Auto 2 Cell Imaging System (ThermoFisher).

The hiPSC-CMs plated on glass coverslips were fixed using the Inside Stain Kit (Miltenyi Biotec) according to manufacturer's instructions. The fixed cells were incubated with α-actinin (1:250, Sigma-Aldrich, #A7811) and myosin heavy chain (1:50, Miltenyi Biotec, #130-112-757) antibodies, followed by Alexa Fluor 594- (1:250, ThermoFisher, #A-21203) and Vio515- (1:100, Miltenyi Biotec, #130-112-760) conjugated secondary antibodies. Nuclei were stained with DAPI and images captured using a confocal laser scanning microscope SP8 (Leica) at 40x magnification.

**Western Blot**

Samples were lysed in RIPA Lysis and Extraction Buffer and total protein measured using the Pierce™ BCA Protein Assay (both Thermo Fisher) according to manufacturer's protocols. The protein samples (40 µg) were loaded on an 8% polyacrylamide gel, and transfer performed using standard protocols[5]. The membrane was incubated with the primary antibodies hERG1a and β-tubulin (1:1000, both Cell Signalling Technology, #12889and#2146, respectively), followed by appropriate HRP-conjugated secondary antibodies (1:10000, Cell Signalling Technology, #7074), and the chemiluminescence signal detected using WesternBright Quantum HRP substrate (Isogen Life Science).

**Patch clamp data acquisition**

Electrophysiological recordings were performed on single hiPSC-CMs. For action potential (AP) and rapid delayed rectifier potassium current ($I_{Kr}$) measurements, spontaneously contracting cardiomyocytes were selected in Tyrode's solution containing (in mM): NaCl 140, KCl 5.4, $CaCl_2$ 1.8, $MgCl_2$ 1.0, glucose 5.5, and HEPES 5.0; pH 7.4 (NaOH). $I_{Kr}$ and APs were measured with the ruptured or perforated patch-clamp technique, respectively, using an Axopatch 200B amplifier (Molecular Devices) at 36 ± 0.2°C. Voltage control and data acquisition of $I_{Kr}$ and APs were performed with pClamp 10.4/Clampfit (Axon Instruments) and custom-made software, respectively. Potentials were corrected for the calculated liquid junction potentials[6], which was 15 mV for both AP and $I_{Kr}$ measurements. Low-resistance patch pipettes (2-3 MΩ; Borosilicate glass capillaries, Harvard Apparatus) were used, and for $I_{Kr}$ measurements, series resistance (Rs) was compensated for ≥80%. $I_{Kr}$ and APs were low-pass filtered (cut-off frequency of 5 kHz and 2 kHz, respectively), and digitised (55 kHz and 40 kHz, respectively).

**Voltage-clamp experiments**

$I_{Kr}$ was measured using 4 s hyper- and depolarising pulses from a holding potential of -40 mV, at a cycle length of 10 s. The extracellular solution was Tyrode's solution, while the pipette solution contained (in mM): K-gluconate 125, KCl 20, $K_2$-ATP 5, HEPES 10, and EGTA 10; pH 7.2 (KOH). The L-type $Ca^{2+}$ current was blocked by adding 5 µM nifedipine (Sigma) to the extracellular solution. $I_{Kr}$ was measured as an E-4031-sensitive current by subtracting the current recorded before and after application of 5 µM E-4031 (Tocris). $I_{Kr}$ density was calculated by dividing current amplitude (pA), measured at the end of the test pulses, by cell membrane capacitance (pF). Cell membrane capacitance was measured by dividing the decay time constant of the capacitive transient in response to 5 mV hyperpolarising steps from -40 mV, by the Rs.

**Current-clamp experiments**

APs were measured in Tyrode's solution, while the pipette solution contained (in mM): K-gluconate 125; KCl 20; NaCl 5.0; amphotericin-B 0.44, and HEPES 10; pH 7.2 (KOH). Because hiPSC-CMs typically have a small or even absent inward rectifying potassium current ($I_{K1}$), the cells have a depolarised resting membrane potential (RMP) and are frequently spontaneously active. We therefore injected an *in silico* 2 pA/pF $I_{K1}$ with kinetics of $Kir_{2.1}$ channels through dynamic clamp, resulting in quiescent hiPSC-CMs with a RMP of less than -75 mV. APs were elicited at 0.2 Hz, 0.5 Hz, 1 Hz, 2 Hz, 3 Hz and 4 Hz by 3 ms, ~1.2x threshold current pulses through the patch pipette. The RMP, upstroke velocity, AP amplitude and AP duration at 20%, 50% and 90% repolarisation were analysed. Parameters from 13 consecutive APs were averaged.

**MEA electrophysiology**

Multi-electrode array (MEA) experiments to record spontaneous electrical activity of hiPSC-CMs were done using 60-electrode MEAs (Multichannel Systems). The MEAs were coated with human fibronectin (40 µg/ml, Alfa Aesar) for 1 h at 37°C, and hiPSC-CMs seeded directly on the electrodes at a density of 2.5 x $10^6$ cells/$cm^2$. Medium was changed at least 1 h before baseline recordings. All measurements were made at 37°C in Medium C and at least 15 min after placing the MEA in the recording system. Extracellular recordings were performed using a MEA1060INV amplifier (Multichannel systems) as previously described[7].

For evaluating E-4031-induced effects on the hiPSC-CMs, sequential addition of increasing concentrations of E-4031 was performed. E-4031 (Tocris) was dissolved in DMSO (Sigma-Aldrich) at 10 mM, with serial dilutions made in Medium C. The final concentration of the drug was achieved by stepwise removal of medium and addition of the same volume of diluted E-

4031 to the well. No more than 7% of the total volume was replaced. The response to each E-4031 concentration was recorded for 1 min after an incubation period of 1 min. The maximum amount of DMSO present in the culture medium was 0.3% at the highest concentration of E-4031. Controls indicated that this percentage of DMSO did not alter the field potential duration (FPD) of the hiPSC-CMs or trigger arrhythmic events.

MEA traces were analysed using Clampfit software to quantify FPD and peak-to-peak intervals over the whole recording time. Bazett's formula for frequency correction was used to correct for FPD dependence on beating rate (cFPD):

$$\frac{FPD}{\sqrt{RR}}$$

where FPD is in ms and RR in s. Arrhythmia-like events were classified and scored based on the following categories: no arrhythmia (0); variable beat period (1); abnormal depolarisations (2); fibrillation (3); quiescent (4).

**Optical evaluation using the Triple Transient Measurement (TTM) system**

The hiPSC-CMs were plated at between 3 – 5.5 x $10^4$ cells per well in black glass-bottom 96-well plates (Greiner), pre-coated with 1:100 Matrigel in DMEM/F-12 (Gibco). Medium was replaced the next day and thereafter every 2-3 days with Pluricyte Cardiomyocyte Medium (PCM; NCardia). Analysis was performed 5-7 days after plating. Cells were incubated for 20 min at 37°C (protected from light) with a voltage sensitive dye (6 µM ANNINE-6plus; Sensitive Farbstoffe), a $Ca^{2+}$ sensitive dye (6 µM Rhod 3; ThermoFisher) and a cell membrane labelling dye (5 µM CellMask Deep Red; ThermoFisher) in PCM for measuring electrical activity, $Ca^{2+}$

flux and contraction, respectively. The dye medium was then removed and PCM added, with cells left to recover for 10-15 min at 37°C protected from light before analysis.

Recording and data processing were performed using a bespoke fast optical switch microscopy system and algorithms developed in-house[8]. Briefly, cells were paced at 1.2 Hz using 10 ms pulses of 14 V with a pair of field stimulation electrodes placed in the culture medium. An Eclipse Ti microscope (Nikon) was fitted with a high-power three LED system (Mightex; 470 nm, 560 nm and 656 nm) including collimators and excitation filters (Semrock; 470±14 nm, 544±12 nm, 650±7.5 nm). A 63x oil immersion objective was used, together with an image intensifier (Photonis) and a high-speed camera (Optronis). Plates were maintained in an environmental chamber at 5% $CO_2$ and 37°C. Three fields of view were measured per well, with recordings of all three fluorescence channels performed at 1000 frames/s for 7 s using fast optical switching of LEDs (1 ms per channel). The three channel signals were then separated and peaks averaged. The 90% durations were then calculated. For voltage this was taken as the top of the peak to 90% of repolarisation, whereas for $Ca^{2+}$ and contraction this was taken from 10% above baseline in the upstroke to 90% of the decay. Five wells were analysed per differentiation of hiPSC-CMs, with three independent differentiations used to generate the mean data per line (n = 15 wells, 45 fields of view).

#### Supplementary Table 1: Sequences of gRNAs and ssODNs used to generate the *KCNH2* variant hiPSC lines

**KCNH2$^{PR/WT}$ clones**

| | |
|---|---|
| **gRNA#1** | 5'-TCGCTACTCAGAGTACGGCG-3' |
| **ssODN_pore** | GGCTGCTGCGGCTGGTGCGCGTGGCGCGGAAGCTGGATCGCTACTCAGAGTACGGCGCGGCCGTGCTGTTCTTGCTCATGTGCACCTTTGCGCTCATC GCGCACTGGCTAGCCTGCATCTGGTACG-3' |

**KCNH2$^{TL/WT}$ clone 1**

| | |
|---|---|
| **gRNA#2** | 5'-TGTAAAAGTCGAAGACCCCC-3' |
| **ssODN_tail** | GACCGGGCGTGGCAGCGGTGGTGCGTCTACCCCGCTCACCCAGCTCTGCTCTCTGGCTGCAGGCGCCTTCTCAGGAGTGTCCATCATTTTCAGCTTCT GGGGGGACAGTCGGGGCCGCCAGTAC-3' |

**KCNH2$^{TL/WT}$ clone 2**

| | |
|---|---|
| **gRNA#3** | 5'-AATGTTGGACACTCCTGAGA-3' |
| **ssODN_tail** | 5'- GACCGGGCGTGGCAGCGGTGGTGCGTCTACCCCGCTCACCCAGCTCTGCTCTCTGGCTGCAGGCGCCTTCTCAGGAGTGTCCATCATTTTCAGCTTCT |
| **ssODN_wt competitor** | 5'- GACCGGGCGTGGCAGCGGTGGTGCGTCGACCCCGCTCACCCAGCTCTGCTCTCTGGCTGCAGGCGCCTTCTCAGGAGTGTCCAACATTTTCAGCTTCT |

#### Supplementary Table 2: List of primers used in this study

| **Primer name** | **Sequence (5' - 3')** | **Purpose** |
|---|---|---|
| **KCNH2_exon 7_Fwd** | CAAGGAGGCAGGTGGTGTAG | amplification of *KCNH2* exon 7 |
| **KCNH2_exon 7_Rev** | CCTCCAACTTGGGTTCCTCC | amplification of *KCNH2* exon 7 |
| **KCNH2_exon 7_seq_Fwd** | CCCCATCAACGGAATGTG | sequencing of *KCNH2* exon 7 |
| **KCNH2_exon 7_seq_Rev** | CACAGCCAATGAGCATGACG | sequencing of *KCNH2* exon 7 |
| **KCNH2_exon 7_cDNA_Fwd** | CTGATCGGGCTGCTGAAGACT | determining *KCNH2* allele transcript expression for $KCNH2^{PR/WT}$ |
| **KCNH2_exon 7_cDNA_Rev** | CCGAAGATGCTAGCGTACATG | determining *KCNH2* allele transcript expression for $KCNH2^{PR/WT}$ |
| **KCNH6_off_target_Fwd** | GCTCTCACTGCTCCTCCATC | amplification of off-target for gRNA#1 |
| **KCNH6_off_target_Rev** | TTCTCGAGTTGGTGTTGGG | amplification of off-target for gRNA#1 |
| **KCNH6_off_target_seq_Fwd** | CTCATCCATGAAGCCTCCCC | sequencing of off-target for gRNA#1 |
| **KCNH6_off_target_seq_Rev** | GCTGAAGGTGAAGTAGAGGG | sequencing of off-target for gRNA#1 |
| **KCNH2_exon 13_Fwd** | CCCTGAGAGCAGTGAGGATG | amplification of *KCNH2* exon 13 |
| **KCNH2_exon 13_Rev** | GGTGGTCACAGCACTGTAGG | amplification of *KCNH2* exon 13 |
| **KCNH2_exon 13_seq_Fwd** | TCAGGTATCCCGGGCGAC | sequencing of *KCNH2* exon 13 |
| **KCNH2_exon 13_seq_Rev** | CTCCCTCTACCAGACAACACC | sequencing of *KCNH2* exon 13 |
| **KCNH2_exon 13_cDNA_Fwd** | CCCTGAGAGCAGTGAGGATG | determining *KCNH2* allele transcript expression for $KCNH2^{TL/WT}$ |
| **KCNH2_exon 13_cDNA_Rev** | GGTGGTCACAGCACTGTAGG | determining *KCNH2* allele transcript expression for $KCNH2^{TL/WT}$ |
| **KCNH2_gRNA#2_IVT_F** | TGTAATACGACTCACTATAGAATGTTGGACACTC CTGAGAGTTTTAGAGCTAGAAATAGC | Forward primer for in vitro transcription of gRNA#2 |
| **KCNH2_gRNA#3_IVT_F** | TGTAATACGACTCACTATAGCCCCCAGAAGCTGA AAATGTGTTTTAGAGCTAGAAATAGC | Forward primer for in vitro transcription of gRNA#3 |
| **gRNA_IVT_Rev** | AGCACCGACTCGGTGCCACT | Reverse primer for in vitro transcription of gRNA#2 and #3 |
| **Off_T.ex13_#2_BEND2_Fwd** | GGGGAAAATTGGGGAAAGGG | assessing potential off-target for gRNA#2 |
| **Off_T.ex13_#2_BEND2_Rev** | CAGCATGTCATGAACTGCAGG | assessing potential off-target for gRNA#2 |
| **Off_T.ex13_#2_HSPA4L_Fwd** | GGTCATGCCCTAAGTCACAGG | assessing potential off-target for gRNA#2 |
| **Off_T.ex13_#2_HSPA4L_Rev** | GACGGGGTTTTGCCATGTTG | assessing potential off-target for gRNA#2 |
| **Off_T.ex13_#2_ETV1_Fwd** | TCCATTTGCGATTTGGTATGGAG | assessing potential off-target for gRNA#2 |
| **Off_T.ex13_#2_ETV1_Rev** | CTGTCTGGCATGTGGGAGTC | assessing potential off-target for gRNA#2 |
| **Off_T.ex13_#2_SGCE_Fwd** | GGTTACCCAGACCGACCTG | assessing potential off-target for gRNA#2 |
| **Off_T.ex13_#2_SGCE_Rev** | GATGTGTTGTTTCCTCCGCC | assessing potential off-target for gRNA#2 |
| **Off_T.ex13_#2_P.20_Fwd** | GTTTGGTCTGAAGCCATGGC | assessing potential off-target for gRNA#2 |
| **Off_T.ex13_#2_P.20_Rev** | CAGGTGGCCTTCGAGGAAAG | assessing potential off-target for gRNA#2 |
| **Off_T.ex13_#3_VWA8_Fwd** | CTTGACTCCCAGCTCTAGCC | assessing potential off-target for gRNA#3 |
| **Off_T.ex13_#3_VWA8_Rev** | CTTTGGGGACTAAGGTGGGG | assessing potential off-target for gRNA#3 |
| **Off_T.ex13_#3_CBL_Fwd** | CCCCTGCTGTGAGACTTCAG | assessing potential off-target for gRNA#3 |
| **Off_T.ex13_#3_CBL_Rev** | AAGGCAGGGGAAAACTGAGG | assessing potential off-target for gRNA#3 |
| **Off_T.ex13_#3_DLX5_Fwd** | GCAAAAACACACACAAGCGC | assessing potential off-target for gRNA#3 |
| **Off_T.ex13_#3_DLX5_Rev** | GCTGTGACCCCCAATCTACC | assessing potential off-target for gRNA#3 |
| **Off_T.ex13_#3_SYT16_Fwd** | CAGTGAGCCTGAAACACAGC | assessing potential off-target for gRNA#3 |
| **Off_T.ex13_#3_SYT16_Rev** | CATGCCCGGCTCTATGCTAG | assessing potential off-target for gRNA#3 |
| **Off_T.ex13_#3_SCN9A_Fwd** | TGTGTCCCCTACCCTGTTCC | assessing potential off-target for gRNA#3 |
| **Off_T.ex13_#3_SCN9A_Rev** | GGTTCAGTACTTTCTTCAGTGGC | assessing potential off-target for gRNA#3 |
| **Off_T.ex13_#3_ACSM5_Fwd** | GTCCATCTGGGGCATCTGAG | assessing potential off-target for gRNA#3 |
| **Off_T.ex13_#3_ACSM5_Rev** | CTCTGCCTCCCGAGTTCAAG | assessing potential off-target for gRNA#3 |

#### Supplementary Table 3: hiPSC lines used in this study

| Cell Line Name | Abbreviation in paper | Reference |
|---|---|---|
| LUMC0020iCTRL-06 | $KCNH2^{WT/WT}$ | wt2 (Zhang, et al 2014) |
| LUMC0020iHERG-01 | $KCNH2^{TL/WT}$ | This study |
| LUMC0020iHERG-02 | $KCNH2^{TL/WT}$ cl2 | This study |
| LUMC0020iHERG-03 | $KCNH2^{PR/WT}$ | This study |
| LUMC0020iHERG-04 | $KCNH2^{PR/WT}$ cl2 | This study |

#### Supplementary Table 4: Coding sequence SNPs and INDELs in genes associated with cardiac arrhythmias and cardiomyopathies in hiPSC line LUMC0020iCTRL04 (KCNH2$^{WT/WT}$)

| Gene | Disease Category | Variant Position | Reference allele | Observed allele | Genotype | RefSNP ID | Transcript | Consequence | allele frequency (1000 Genomes) | Predicted pathogenicity |
|---|---|---|---|---|---|---|---|---|---|---|
| *ABCC9* | arrhythmia; cardiomyopathy | chr12_22063115 | A | G | hom | rs10770865 | NM_005691.3:p.Pro432Pro/c.1296T>C | synonymous_variant | 1.00 | benign |
| *ACTA1* | cardiomyopathy | none | | | | | | | | |
| *ACTC1* | cardiomyopathy | none | | | | | | | | |
| *ACTN2* | cardiomyopathy | chr1_236882303 | T | C | hom | rs1341864 | NM_001103.3:p.Ile117Ile/c.351T>C | synonymous_variant | 0.99 | benign |
| | | chr1_236883421 | C | T | hom | rs1341863 | NM_001103.3:p.Asn126Asn/c.378C>T | synonymous_variant | 0.92 | benign |
| | | chr1_236925844 | G | A | het | rs12063382 | NM_001103.3:p.Ser870Ser/c.2610G>A | synonymous_variant | 0.20 | benign |
| | | chr1_236899899 | TC | T | hom | rs11355106 | NM_001278344.1:p.Pro32fs/c.95delC | frameshift_variant | 0.53 | likely benign |
| *AKAP9* | arrhythmia | chr7_91630620 | G | T | het | rs6964587 | NM_005751.4:p.Met463Ile/c.1389G>T | missense_variant | 0.37 | benign |
| | | chr7_91712698 | A | G | het | rs6960867 | NM_005751.4:p.Asn2792Ser/c.8375A>G | missense_variant | 0.30 | benign |
| | | chr7_91714911 | C | T | hom | rs1063242 | NM_005751.4:p.Pro2979Ser/c.8935C>T | missense_variant | 1.00 | benign |
| | | chr7_91632306 | C | T | hom | rs1989779 | NM_005751.4:p.Thr1025Thr/c.3075C>T | synonymous_variant | 0.94 | benign |
| | | chr7_91641928 | A | G | het | rs13245393 | NM_005751.4:p.Glu1168Glu/c.3504A>G | synonymous_variant | 0.37 | benign |
| | | chr7_91691601 | C | T | het | rs10236397 | NM_005751.4:p.Gly1926Gly/c.5778C>T | synonymous_variant | 0.36 | benign |
| | | chr7_91713972 | C | T | het | rs10228334 | NM_005751.4:p.Leu2889Leu/c.8665C>T | synonymous_variant | 0.37 | benign |
| | | chr7_91715662 | C | T | het | rs28927678 | NM_005751.4:p.Leu3049Leu/c.9145C>T | synonymous_variant | 0.30 | benign |
| | | chr7_91726927 | A | C | het | rs1063243 | NM_005751.4:p.Arg3476Arg/c.10426A>C | synonymous_variant | 0.38 | benign |
| | | chr7_91652178 | A | AAAC | het | rs10644111 | NM_005751.4:p.Lys1335_Leu1336insGln/c.4004_4006dupAAC | inframe_insertion | 0.42 | benign |
| *ANK2* | arrhythmia | none | | | | | | | | |
| *ANKRD1* | cardiomyopathy | none | | | | | | | | |
| *BAG3* | cardiomyopathy | chr10_121436286 | C | T | het | rs3858340 | NM_004281.3:p.Pro407Leu/c.1220C>T | missense_variant | 0.16 | benign |
| | | chr10_121436068 | T | G | het | rs3858339 | NM_004281.3:p.Pro334Pro/c.1002T>G | synonymous_variant | 0.16 | benign |
| | | chr10_121436362 | A | G | hom | rs196295 | NM_004281.3:p.Val432Val/c.1296A>G | synonymous_variant | 0.71 | benign |
| *BRAF* | cardiomyopathy | none | | | | | | | | |
| *CACNA1C* | arrhythmia; cardiomyopathy | chr12_2791130 | C | T | hom | rs10848683 | NM_199460.3:p.Pro1868Leu/c.5603C>T | missense_variant | 0.67 | benign |
| | | chr12_2791132 | A | G | hom | rs10774053 | NM_199460.3:p.Met1869Val/c.5605A>G | missense_variant | 0.77 | benign |
| | | chr12_2721137 | C | T | het | rs216008 | NM_199460.3:p.Phe1282Phe/c.3846C>T | synonymous_variant | 0.26 | benign |
| *CACNA2D1* | arrhythmia | chr7_81588636 | G | A | hom | rs1229502 | NM_000722.3:p.Pro1038Pro/c.3114C>T | synonymous_variant | 0.22 | benign |
| *CACNB2* | arrhythmia | chr10_18828635 | T | G | het | rs58225473 | NM_201596.2:p.Asp655Glu/c.1965T>G | missense_variant | 0.10 | benign |
| *CALM1* | arrhythmia | none | | | | | | | | |
| *CALM2* | arrhythmia | none | | | | | | | | |
| *CALM3* | arrhythmia | none | | | | | | | | |
| *CALR3* | cardiomyopathy | chr19_16591464 | G | A | het | rs9305079 | NM_145046.4:p.Tyr324Tyr/c.972C>T | synonymous_variant | 0.65 | benign |
| | | chr19_16601194 | C | T | het | rs3810198 | NM_145046.4:p.Gln127Gln/c.381G>A | synonymous_variant | 0.66 | benign |
| *CASQ2* | arrhythmia; cardiomyopathy | none | | | | | | | | |
| *CAV3* | arrhythmia; cardiomyopathy | chr3_8775661 | C | T | het | rs1008642 | NM_001234.4:p.Asn33Asn/c.99C>T | synonymous_variant | 0.37 | benign |
| | | chr3_8787220 | T | C | hom | rs13087941 | NM_001234.4:p.Phe41Phe/c.123T>C | synonymous_variant | 0.15 | benign |
| *CBL* | cardiomyopathy | none | | | | | | | | |
| *COX15* | cardiomyopathy | chr10_101473218 | A | G | hom | rs2231687 | NM_004376.6:p.Phe374Leu/c.1120T>C | missense_variant | 0.83 | benign |
| *CRYAB* | cardiomyopathy | none | | | | | | | | |
| *CSRP3* | cardiomyopathy | none | | | | | | | | |
| *DES* | cardiomyopathy | chr2_220283259 | A | G | hom | rs1318299 | NM_001927.3:p.Pro25Pro/c.75A>G | synonymous_variant | 0.89 | benign |
| | | chr2_220283277 | T | C | hom | rs2017800 | NM_001927.3:p.Ser31Ser/c.93T>C | synonymous_variant | 0.87 | benign |
| | | chr2_220285309 | C | T | hom | rs1058261 | NM_001927.3:p.Asp276Asp/c.828C>T | synonymous_variant | 0.34 | benign |

| | | | | | | | | | | |
|---|---|---|---|---|---|---|---|---|---|---|
| | | chr2_220285666 | G | C | hom | rs12920 | NM_001927.3:p.Leu338Leu/c.1014G>C | synonymous_variant | 0.34 | benign |
| | | chr2_220286142 | G | A | hom | rs1058284 | NM_001927.3:p.Ala368Ala/c.1104G>A | synonymous_variant | 0.33 | benign |
| *DMD* | cardiomyopathy | chrX_31496350 | C | T | hom | rs1800280 | NM_004006.2:p.Arg2937Gln/c.8810G>A | missense_variant | 0.88 | benign |
| | | chrX_31893307 | T | G | het | rs1800275 | NM_004006.2:c.7095+1A>C | splice_donor_variant+intron_variant | 0.18 | benign |
| | | chrX_32486756 | C | T | het | rs1800268 | NM_004006.2:p.Ser1007Ser/c.3021G>A | synonymous_variant | 0.01 | benign |
| *DNAJC19* | cardiomyopathy | none | | | | | | | | |
| *DOLK* | cardiomyopathy | chr9_131708952 | G | A | het | rs145310298 | NM_014908.3:p.Arg211Cys/c.631C>T | missense_variant | 0.00 | likely benign; uncertain signficance |
| *DSC2* | cardiomyopathy | chr18_28649042 | T | C | hom | rs1893963 | NM_024422.4:p.Ile776Val/c.2326A>G | missense_variant | 0.20 | benign |
| *DSG2* | cardiomyopathy | none | | | | | | | | |
| *DSP* | cardiomyopathy | chr6_7581636 | G | A | het | rs6929069 | NM_004415.3:p.Arg1738Gln/c.5213G>A | missense_variant | 0.24 | benign |
| | | chr6_7563983 | T | G | hom | rs2806234 | NM_004415.3:p.Ala247Ala/c.741T>G | synonymous_variant | 1.00 | benign |
| | | chr6_7577260 | C | T | het | rs2064217 | NM_004415.3:p.Cys954Cys/c.2862C>T | synonymous_variant | 0.28 | benign |
| *DTNA* | cardiomyopathy | chr18_32470291 | G | A | hom | rs9944927 | NM_001198938.1:p.Glu688Lys/c.2062G>A | missense_variant | 0.27 | benign |
| *EMD* | cardiomyopathy | none | | | | | | | | |
| *FHL1* | cardiomyopathy | none | | | | | | | | |
| *FKTN* | cardiomyopathy | chr9_108366734 | G | A | het | rs34787999 | NM_001079802.1:p.Arg203Gln/c.608G>A | missense_variant | 0.16 | benign |
| | | chr9_108380355 | C | A | het | rs17309806 | NM_001079802.1:p.Leu342Leu/c.1026C>A | synonymous_variant | 0.18 | benign |
| *FXN* | cardiomyopathy | chr9_71650752 | A | G | hom | rs2481598 | NM_000144.4:p.Pro18Pro/c.54A>G | synonymous_variant | 0.98 | benign |
| *GAA* | cardiomyopathy | chr17_78079597 | A | G | hom | rs1042393 | NM_000152.4:p.His199Arg/c.596A>G | missense_variant | 0.60 | benign |
| | | chr17_78079669 | G | A | hom | rs1042395 | NM_000152.4:p.Arg223His/c.668G>A | missense_variant | 0.60 | benign |
| | | chr17_78091405 | G | A | hom | rs1126690 | NM_000152.4:p.Val780Ile/c.2338G>A | missense_variant | 0.71 | benign |
| | | chr17_78078709 | T | C | hom | rs1800300 | NM_000152.4:p.Cys108Cys/c.324T>C | synonymous_variant | 0.71 | benign |
| | | chr17_78079643 | C | T | het | rs1800301 | NM_000152.4:p.Ser214Ser/c.642C>T | synonymous_variant | 0.11 | benign |
| | | chr17_78082504 | G | A | hom | rs1800304 | NM_000152.4:p.Gln401Gln/c.1203G>A | synonymous_variant | 0.60 | benign |
| | | chr17_78084769 | G | A | het | rs1042396 | NM_000152.4:p.Arg527Arg/c.1581G>A | synonymous_variant | 0.16 | benign |
| | | chr17_78087109 | A | G | het | rs1800310 | NM_000152.4:p.Thr711Thr/c.2133A>G | synonymous_variant | 0.24 | benign |
| | | chr17_78092063 | G | A | hom | rs1042397 | NM_000152.4:p.Gly851Gly/c.2553G>A | synonymous_variant | 0.51 | benign |
| *GATAD1* | cardiomyopathy | none | | | | | | | | |
| *GLA* | cardiomyopathy | none | | | | | | | | |
| *GPD1L* | arrhythmia | chr3_32181761 | C | T | het | rs9835387 | NM_015141.3:p.Asp136Asp/c.408C>T | synonymous_variant | 0.15 | benign |
| *HCN4* | arrhythmia | chr15_73614834 | T | C | hom | rs529004 | NM_005477.2:p.Pro1200Pro/c.3600A>G | synonymous_variant | 0.86 | benign |
| *ILK* | cardiomyopathy | none | | | | | | | | |
| *JPH2* | cardiomyopathy | chr20_42743454 | A | G | het | rs6093935 | NM_020433.4:p.Phe691Phe/c.2073T>C | synonymous_variant | 0.26 | benign |
| | | chr20_42744587 | G | C | het | rs74352869 | NM_020433.4:p.Pro576Pro/c.1728C>G | synonymous_variant | 0.15 | benign |
| | | chr20_42815190 | G | A | hom | rs1883790 | NM_020433.4:p.Tyr52Tyr/c.156C>T | synonymous_variant | 0.85 | benign |
| *JUP* | cardiomyopathy | chr17_39912145 | T | A | het | rs1126821 | NM_002230.2:p.Met697Leu/c.2089A>T | missense_variant | 0.59 | benign |
| | | chr17_39925925 | A | G | hom | rs7405731 | NM_002230.2:p.Asp71Asp/c.213T>C | synonymous_variant | 0.72 | benign |
| *KCND3* | arrhythmia | chr1_112524680 | C | G | het | rs17215423 | NM_004980.4:p.Ser223Ser/c.669G>C | synonymous_variant | 0.01 | benign |
| *KCNE1* | arrhythmia | chr21_35821821 | T | C | het | rs1805127 | NM_001330065.1:p.Ser41Gly/c.121A>G | missense_variant | 0.67 | benign |
| *KCNE2* | arrhythmia | none | | | | | | | | |
| *KCNE3* | arrhythmia | none | | | | | | | | |
| *KCNH2* | arrhythmia | chr7_150645534 | T | G | het | rs1805123 | NM_000238.3:p.Lys897Thr/c.2690A>C | missense_variant +splice_region_variant | 0.14 | benign |
| | | chr7_150648198 | A | G | hom | rs1137617 | NM_000238.3:p.Tyr652Tyr/c.1956T>C | synonymous_variant | 0.77 | benign |
| | | chr7_150648789 | T | C | het | rs1805121 | NM_000238.3:p.Leu564Leu/c.1692A>G | synonymous_variant | 0.61 | benign |
| | | chr7_150649531 | G | A | het | rs1805120 | NM_000238.3:p.Phe513Phe/c.1539C>T | synonymous_variant | 0.34 | benign |
| | | chr7_150649603 | G | A | het | rs740952 | NM_000238.3:p.Ile489Ile/c.1467C>T | synonymous_variant | 0.34 | benign |
| *KCNJ2* | arrhythmia | none | | | | | | | | |

| | | | | | | | | | | |
|---|---|---|---|---|---|---|---|---|---|---|
| *KCNJ5* | arrhythmia | chr11_128782012 | C | G | hom | rs7102584 | NM_000890.3:p.Gln282Glu/c.844C>G | missense_variant | 1.00 | benign |
| | | chr11_128781339 | T | C | hom | rs6590357 | NM_000890.3:p.Ser57Ser/c.171T>C | synonymous_variant | 0.87 | benign |
| | | chr11_128781978 | T | G | hom | rs7118824 | NM_000890.3:p.Leu270Leu/c.810T>G | synonymous_variant | 0.87 | benign |
| | | chr11_128782002 | T | C | hom | rs7118833 | NM_000890.3:p.His278His/c.834T>C | synonymous_variant | 0.87 | benign |
| *KCNJ8* | arrhythmia | none | | | | | | | | |
| *KCNQ1* | arrhythmia; cardiomyopathy | none | | | | | | | | |
| *KLF10* | cardiomyopathy | none | | | | | | | | |
| *KRAS* | cardiomyopathy | chr12_25368462 | C | T | hom | rs4362222 | NM_033360.3:p.Arg161Arg/c.483G>A | synonymous_variant | 1.00 | benign |
| *LAMA2* | cardiomyopathy | chr6_129807629 | C | T | het | rs2229848 | NM_000426.3:p.Ala2587Val/c.7760C>T | missense_variant | 0.58 | benign |
| | | chr6_129371106 | C | T | het | rs1140366 | NM_000426.3:p.Ile52Ile/c.156C>T | synonymous_variant | 0.07 | benign |
| | | chr6_129381026 | C | A | hom | rs4404787 | NM_000426.3:p.Thr127Thr/c.381C>A | synonymous_variant | 0.94 | benign |
| | | chr6_129612808 | A | G | het | rs1027199 | NM_000426.3:p.Gln933Gln/c.2799A>G | synonymous_variant | 0.31 | benign |
| | | chr6_129762112 | G | A | het | rs2297738 | NM_000426.3:p.Thr2079Thr/c.6237G>A | synonymous_variant | 0.18 | benign |
| | | chr6_129807699 | G | C | het | rs2229849 | NM_000426.3:p.Val2610Val/c.7830G>C | synonymous_variant | 0.58 | benign |
| | | chr6_129807714 | G | A | het | rs2229850 | NM_000426.3:p.Pro2615Pro/c.7845G>A | synonymous_variant | 0.40 | benign |
| | | chr6_129813175 | T | C | het | rs35313209 | NM_000426.3:p.Asn2676Asn/c.8028T>C | synonymous_variant | 0.02 | benign |
| *LAMA4* | cardiomyopathy | chr6_112457390 | C | T | het | rs2032567 | NM_001105206.2:p.Gly1117Ser/c.3349G>A | missense_variant | 0.84 | benign |
| | | chr6_112493872 | A | G | hom | rs1050348 | NM_001105206.2:p.Tyr498His/c.1492T>C | missense_variant | 0.76 | benign |
| | | chr6_112508769 | T | G | hom | rs9387061 | NM_001105206.2:p.Ala283Ala/c.849A>C | synonymous_variant | 1.00 | benign |
| | | chr6_112512905 | G | A | het | rs2072021 | NM_001105206.2:p.Thr217Thr/c.651C>T | synonymous_variant | 0.32 | benign |
| *LAMP2* | cardiomyopathy | none | | | | | | | | |
| *LDB3* | cardiomyopathy | none | | | | | | | | |
| *LMNA* | cardiomyopathy | none | | | | | | | | |
| *MAP2K1* | cardiomyopathy | none | | | | | | | | |
| *MAP2K2* | cardiomyopathy | none | | | | | | | | |
| *MIB1* | cardiomyopathy | none | | | | | | | | |
| *MURC* | cardiomyopathy | chr9_103348634 | G | A | het | rs2780956 | NM_001018116.2:p.Arg332Arg/c.996G>A | synonymous_variant | 0.29 | benign |
| *MYBPC3* | cardiomyopathy | chr11_47371598 | C | T | het | rs3729986 | NM_000256.3:p.Val158Met/c.472G>A | missense_variant | 0.03 | benign |
| *MYH6* | cardiomyopathy | chr14_23861811 | A | G | het | rs365990 | NM_002471.3:p.Val1101Ala/c.3302T>C | missense_variant | 0.37 | benign |
| | | chr14_23855569 | A | G | het | rs178640 | NM_002471.3:p.Ala1638Ala/c.4914T>C | synonymous_variant | 0.49 | benign |
| | | chr14_23862710 | C | T | het | rs145274612 | NM_002471.3:p.Glu982Glu/c.2946G>A | synonymous_variant | 0.00 | benign |
| | | chr14_23874507 | G | T | het | rs2277473 | NM_002471.3:p.Arg143Arg/c.427C>A | synonymous_variant | 0.08 | benign |
| | | chr14_23874523 | C | T | het | rs2277474 | NM_002471.3:p.Glu137Glu/c.411G>A | synonymous_variant | 0.15 | benign |
| *MYH7* | cardiomyopathy | chr14_23892888 | A | G | het | rs7157716 | NM_000257.3:p.Ile989Ile/c.2967T>C | synonymous_variant | 0.38 | benign |
| | | chr14_23899060 | G | A | het | rs735712 | NM_000257.3:p.Gly354Gly/c.1062C>T | synonymous_variant | 0.06 | benign |
| | | chr14_23902753 | G | A | het | rs2069540 | NM_000257.3:p.Thr63Thr/c.189C>T | synonymous_variant | 0.52 | benign |
| *MYL2* | cardiomyopathy | chr12_111353556 | A | G | het | rs2301610 | NM_000432.3:p.Ile44Ile/c.132T>C | synonymous_variant | 0.12 | benign |
| *MYL3* | cardiomyopathy | none | | | | | | | | |
| *MYLK2* | cardiomyopathy | chr20_30409452 | T | C | het | rs6058469 | NM_033118.3:p.Ile228Ile/c.684T>C | synonymous_variant | 0.07 | benign |
| *MYO6* | cardiomyopathy | none | | | | | | | | |
| *MYOZ2* | cardiomyopathy | none | | | | | | | | |
| *MYPN* | cardiomyopathy | chr10_69926334 | C | G | het | rs10823148 | NM_001256267.1:p.Phe628Leu/c.1884C>G | missense_variant | 0.32 | benign |
| | | chr10_69933921 | G | A | het | rs10997975 | NM_001256267.1:p.Ser691Asn/c.2072G>A | missense_variant | 0.34 | benign |
| | | chr10_69933969 | G | A | het | rs7916821 | NM_001256267.1:p.Ser707Asn/c.2120G>A | missense_variant | 0.33 | benign |
| | | chr10_69934258 | C | G | het | rs3814182 | NM_001256267.1:p.Ser803Arg/c.2409C>G | missense_variant | 0.47 | benign |
| | | chr10_69959242 | C | A | het | rs7079481 | NM_001256267.1:p.Pro1135Thr/c.3403C>A | missense_variant | 0.34 | benign |
| | | chr10_69909802 | G | A | het | rs10997948 | NM_001256267.1:p.Gln417Gln/c.1251G>A | synonymous_variant | 0.08 | benign |
| | | chr10_69926097 | T | C | het | rs2673794 | NM_001256267.1:p.Ser549Ser/c.1647T>C | synonymous_variant | 0.48 | benign |
| | | chr10_69948844 | T | C | hom | rs10733838 | NM_001256267.1:p.Val962Val/c.2886T>C | synonymous_variant | 0.98 | benign |
| *NEBL* | cardiomyopathy | chr10_21108377 | C | T | het | rs1006363 | NM_006393.2:p.Arg677Arg/c.2031G>A | synonymous_variant | 0.18 | benign |

| | | | | | | | | | | |
|---|---|---|---|---|---|---|---|---|---|---|
| *NEXN* | cardiomyopathy | chr1_78392446 | G | A | het | rs1166698 | NM_144573.3:p.Gly245Arg/c.733G>A | missense_variant | 0.15 | benign |
| *NPPA* | cardiomyopathy | none | | | | | | | | |
| *NRAS* | cardiomyopathy | none | | | | | | | | |
| *PDLIM3* | cardiomyopathy | none | | | | | | | | |
| *PKP2* | arrhythmia; cardiomyopathy | none | | | | | | | | |
| *PLN* | cardiomyopathy | none | | | | | | | | |
| *PRDM16* | cardiomyopathy | chr1_3328358 | T | C | hom | rs870124 | NM_022114.3:p.Ser533Pro/c.1597T>C | missense_variant | 0.95 | benign |
| | | chr1_3328659 | C | T | het | rs2493292 | NM_022114.3:p.Pro633Leu/c.1898C>T | missense_variant | 0.11 | benign |
| | | chr1_3301721 | C | T | het | rs2282198 | NM_022114.3:p.Ser148Ser/c.444C>T | synonymous_variant | 0.34 | benign |
| *PRKAG2* | cardiomyopathy | none | | | | | | | | |
| *PTPN11* | cardiomyopathy | none | | | | | | | | |
| *RAF1* | cardiomyopathy | none | | | | | | | | |
| *RBM20* | cardiomyopathy | chr10_112595719 | G | C | hom | rs942077 | NM_001134363.2:p.Glu1223Gln/c.3667G>C | missense_variant | 0.70 | benign |
| | | chr10_112404302 | G | A | het | rs35141404 | NM_001134363.2:p.Arg30Arg/c.90G>A | synonymous_variant | 0.22 | benign |
| *RANGRF* | arrhythmia | none | | | | | | | | |
| *RYR2* | arrhythmia; cardiomyopathy | chr1_237841390 | A | G | het | rs34967813 | NM_001035.2:p.Gln2958Arg/c.8873A>G | missense_variant | 0.10 | benign |
| | | chr1_237617757 | C | T | het | rs3765097 | NM_001035.2:p.Ser453Ser/c.1359C>T | synonymous_variant | 0.54 | benign |
| | | chr1_237711797 | A | G | hom | rs2253273 | NM_001035.2:p.Ser991Ser/c.2973A>G | synonymous_variant | 0.83 | benign |
| | | chr1_237801770 | T | C | hom | rs707189 | NM_001035.2:p.Leu2302Leu/c.6906T>C | synonymous_variant | 0.95 | benign |
| | | chr1_237814783 | C | T | hom | rs684923 | NM_001035.2:p.His2602His/c.7806C>T | synonymous_variant | 0.55 | benign |
| | | chr1_237863718 | T | G | hom | rs2797436 | NM_001035.2:p.Ser3106Ser/c.9318T>G | synonymous_variant | 0.97 | benign |
| | | chr1_237881770 | C | T | hom | rs2797441 | NM_001035.2:p.Thr3501Thr/c.10503C>T | synonymous_variant | 0.96 | benign |
| | | chr1_237890437 | C | T | hom | rs2685301 | NM_001035.2:p.Ser3592Ser/c.10776C>T | synonymous_variant | 0.96 | benign |
| *SCN3B* | arrhythmia | none | | | | | | | | |
| *SCN4B* | arrhythmia | none | | | | | | | | |
| *SCN5A* | arrhythmia; cardiomyopathy | chr3_38592406 | A | G | het | rs1805126 | NM_001099404.1:p.Asp1819Asp/c.5457T>C | synonymous_variant | 0.49 | benign |
| | | chr3_38622467 | T | C | hom | rs7430407 | NM_001099404.1:p.Glu1061Glu/c.3183A>G | synonymous_variant | 0.92 | benign |
| | | chr3_38674712 | T | C | hom | rs6599230 | NM_001099404.1:p.Ala29Ala/c.87A>G | synonymous_variant | 0.78 | benign |
| *SGCB* | cardiomyopathy | none | | | | | | | | |
| *SGCD* | cardiomyopathy | chr5_155771579 | T | C | het | rs1801193 | NM_000337.5:p.Tyr28Tyr/c.84T>C | synonymous_variant | 0.49 | benign |
| *SHOC2* | cardiomyopathy | none | | | | | | | | |
| *SLC25A4* | cardiomyopathy | none | | | | | | | | |
| *SNTA1* | arrhythmia | none | | | | | | | | |
| *TAZ* | cardiomyopathy | none | | | | | | | | |
| *TCAP* | cardiomyopathy | chr17_37822311 | A | C | hom | rs1053651 | NM_003673.3:p.Ala151Ala/c.453A>C | synonymous_variant | 0.55 | benign |
| *TGFB3* | cardiomyopathy | none | | | | | | | | |
| *TMEM43* | cardiomyopathy | chr3_14174427 | A | T | het | rs4685076 | NM_024334.2:p.Lys168Asn/c.504A>T | missense_variant | 0.35 | benign |
| | | chr3_14175262 | T | C | het | rs2340917 | NM_024334.2:p.Met179Thr/c.536T>C | missense_variant | 0.46 | benign |
| *TMPO* | cardiomyopathy | chr12_98927830 | C | G | het | rs17459334 | NM_003276.2:p.Gln599Glu/c.1795C>G | missense_variant | 0.06 | benign |
| *TNNC1* | cardiomyopathy | none | | | | | | | | |
| *TNNI3* | cardiomyopathy | none | | | | | | | | |
| *TNNT2* | cardiomyopathy | chr1_201334382 | G | A | hom | rs3729547 | NM_001276345.1:p.Ile116Ile/c.348C>T | synonymous_variant | 0.70 | benign |
| *TPM1* | cardiomyopathy | chr15_63335949 | G | A | het | rs1312883932 | NM_001301244.1:p.Ser53Ser/c.159G>A | synonymous_variant | 0.00 | benign |
| | | chr15_63351840 | C | A | hom | rs1071646 | NM_000366.5:p.Ala151Ala/c.453C>A | synonymous_variant | 0.71 | benign |
| *TRDN* | arrhythmia | chr6_123687288 | A | C | hom | rs2873479 | NM_006073.3:p.Ile438Ser/c.1313T>G | missense_variant | 0.94 | benign |
| | | chr6_123833457 | G | C | hom | rs6902416 | NM_006073.3:p.Leu201Val/c.601C>G | missense_variant | 0.84 | benign |
| | | chr6_123869607 | G | C | hom | rs9490809 | NM_006073.3:p.Thr128Ser/c.383C>G | missense_variant | 0.39 | benign |
| *TRIM63* | cardiomyopathy | none | | | | | | | | |
| *TTN* | cardiomyopathy | chr2_179444768 | C | G | hom | rs4145333 | NM_001267550.2:p.Ala22416Pro/c.67246G>( | missense_variant | 0.99 | benign |
| | | chr2_179612383 | C | T | het | rs72648907 | NM_133379.4:p.Arg4915His/c.14744G>A | missense_variant | 0.01 | benign |

| | | | | | | | | | | |
|---|---|---|---|---|---|---|---|---|---|---|
| | | chr2_179615887 | T | C | hom | rs922984 | NM_133379.4:p.Asp3747Gly/c.11240A>G | missense_variant | 0.73 | benign |
| | | chr2_179615931 | C | G | hom | rs922985 | NM_133379.4:p.Leu3732Phe/c.11196G>C | missense_variant | 0.98 | benign |
| | | chr2_179623758 | C | T | hom | rs2291310 | NM_001267550.2:p.Ser3419Asn/c.10256G>A | missense_variant | 0.81 | benign |
| | | chr2_179629461 | C | T | hom | rs2291311 | NM_001267550.2:p.Val3261Met/c.9781G>A | missense_variant | 0.80 | benign |
| | | chr2_179644035 | G | A | hom | rs1552280 | NM_001267550.2:p.Ser1295Leu/c.3884C>T | missense_variant | 0.92 | benign |
| | | chr2_179644855 | T | C | hom | rs10497520 | NM_001267550.2:p.Lys1201Glu/c.3601A>G | missense_variant | 0.50 | benign |
| | | chr2_179650408 | G | A | hom | rs35813871 | NM_001267550.2:p.Thr811Ile/c.2432C>T | missense_variant | 0.10 | benign |
| | | chr2_179620951 | C | T | hom | rs7585334 | NM_001267550.2:p.Gly3751Asp/c.11252G>A | missense_variant | 0.80 | benign |
| | | chr2_179585266 | C | T | hom | rs2562831 | NM_001267550.2:p.Gln7741Gln/c.23223G>A | synonymous_variant | 0.98 | benign |
| | | chr2_179600563 | G | A | hom | rs2742348 | NM_001267550.2:p.Ser4870Ser/c.14610C>T | synonymous_variant | 0.98 | benign |
| | | chr2_179614952 | A | G | hom | rs10803917 | NM_133379.4:p.Leu4059Leu/c.12175T>C | synonymous_variant | 0.98 | benign |
| | | chr2_179615994 | T | C | hom | rs922986 | NM_133379.4:p.Thr3711Thr/c.11133A>G | synonymous_variant | 0.98 | benign |
| | | chr2_179629363 | T | C | hom | rs4894043 | NM_001267550.2:p.Glu3293Glu/c.9879A>G | synonymous_variant | 0.97 | benign |
| *TXNRD2* | cardiomyopathy | chr22_19868218 | A | G | het | rs1139793 | NM_006440.4:p.Ile370Thr/c.1109T>C | missense_variant | 0.72 | benign |
| | | chr22_19882984 | T | G | het | rs5992495 | NM_006440.4:p.Ser299Arg/c.895A>C | missense_variant | 0.25 | benign |
| | | chr22_19907099 | C | A | het | rs5748469 | NM_006440.4:p.Ala66Ser/c.196G>T | missense_variant | 0.48 | benign |
| | | chr22_19867771 | C | T | het | rs1139795 | NM_006440.4:p.Pro402Pro/c.1206G>A | synonymous_variant | 0.27 | benign |
| | | chr22_19906511 | G | A | het | rs11541479 | NM_006440.4:p.Leu82Leu/c.246C>T | synonymous_variant | 0.17 | benign |
| | | chr22_19907118 | G | A | hom | rs5748470 | NM_006440.4:p.Ala59Ala/c.177C>T | synonymous_variant | 0.60 | benign |
| *VCL* | cardiomyopathy | chr10_75865065 | G | A | hom | rs767809 | NM_014000.2:p.Pro796Pro/c.2388G>A | synonymous_variant | 0.32 | benign |
| | | chr10_75871735 | C | G | hom | rs2131956 | NM_014000.2:p.Gly938Gly/c.2814C>G | synonymous_variant | 0.62 | benign |

#### Supplementary Table 5: Predicted gRNA off-target exonic sequences analysed for each *KCNH2* variant hiPSC line

| | Off-target Sequence | Mismatch Position | Mismatch | Score | Chromosome | Strand | Gene |
|---|---|---|---|---|---|---|---|
| **$KCNH2^{PR/WT}$ clones** | CCGCTACTCTGAGTATGGGGCGG | *........*.....*..*. | 4 | 0.02 | chr17 | + | ***KCNH6*** |
| **$KCNH2^{TL/WT}$ clone 1** | CTACCAGAAAATGAAAATGTGGG | .**......**......... | 4 | 0.55 | chrX | - | ***BEND2*** |
| | TCCCCAGAAGCAAAAAATTTAGG | *..........**.....*. | 4 | 0.49 | chr4 | - | ***HSPA4L*** |
| | AGCCCAGAAACTGAAAATATTGG | **.......*........*. | 4 | 0.46 | chr7 | - | ***ETV1*** |
| | TCCCCAACAGCTGAAAATGTGGG | *.....**............ | 3 | 0.42 | chr7 | - | ***SGCE*** |
| | CTGTAAGAAGCTGAAAATGTTGG | .****............... | 4 | 0.2 | chr1 | + | ***PRAMEF20*** |
| **$KCNH2^{TL/WT}$ clone 2** | AATGCAGGACACTTCAGAGAAGG | ....**.......*.*.... | 4 | 0.58 | chr16 | - | **ACSM5** |
| | AAAGTTGGACATTCCAAAGATGG | ..*........*...**... | 4 | 0.33 | chr2 | + | ***SCN9A*** |
| | AATGATGGATAATCCTGAAATGG | ....*....*.*......*. | 4 | 0.24 | chr14 | + | ***SYT16*** |
| | AAAGTTGGCGATTCCTGAGACGG | ..*.....**.*........ | 4 | 0.09 | chr7 | + | ***DLX5*** |
| | TAGGATGGACTCTCCTGAGAAGG | *.*.*.....*......... | 4 | 0.07 | chr11 | + | ***CBL*** |
